# Diffusion MRI Processing in the HEALthy Brain and Child Development Study: Innovations and Applications

**DOI:** 10.1101/2025.11.10.687672

**Authors:** Matthew Cieslak, M. Okan Irfanoglu, Steven L. Meisler, Taylor Salo, Adam Raikes, Philip A Cook, Ai Wern Chung, Erik G. Lee, Ruolin Li, Xu Li, Diliana Pecheva, Damien A. Fair, Christopher D. Smyser, Michael P. Harms, Bennett A. Landman, Jessica Wisnowski, Hao Huang, Andrew L. Alexander, Theodore D. Satterthwaite

## Abstract

The landmark ongoing HEALthy Brain and Cognitive Development (HBCD) study will longitudinally chart brain development in a large sample (projected *n*=7,200) of infants through age 10 years with multimodal neuroimaging that includes an advanced diffusion MRI (dMRI) acquisition. Here, we detail advances in dMRI image processing developed for HBCD, incorporated into the widely used *QSIPrep* pipeline. Major changes to preprocessing include improvements in infant brain extraction, distortion correction, and normalization to infant-specific templates. Additionally, we describe a new software package – *QSIRecon* – that yields rich derived data including diverse maps of tissue microstructure as well as person-specific white matter bundles. Using dMRI data from a subset of the HBCD 1.0 release where age information was available (*n*=529 sessions across two time points), we observe critical improvements in data quality with preprocessing and see expected developmental patterns. Moving forward, the publicly-available data from HBCD will rapidly grow to become the largest study of brain development in infancy and early childhood using dMRI. *QSIPrep* and *QSIRecon* are openly available and can be applied to other infant and pediatric dMRI datasets.

## INTRODUCTION

During infancy, white matter development plays a central role in the emergence of motor, cognitive, emotional, and behavioral functions (Filley & Fields, 2016; Kim et al., 2025; Lebel & Deoni, 2018; Schilling et al., 2023; Ouyang et al., 2019). Studying the specific trajectories of white matter maturation during this period can provide foundational insight into normative brain development and help identify early markers of risk for neurodevelopmental and psychiatric disorders (Fields, 2008; Spann & Rogers, 2023). Diffusion-weighted MRI (dMRI) is the primary non-invasive technique to study white matter across the human lifespan. The ongoing multi-site HEALthy Brain and Child Development (HBCD) study is the largest study of early brain development (projected sample size: *n*=7,200) and includes an advanced dMRI acquisition (Dean et al., 2024). Here we describe advances in dMRI image processing specifically developed for HBCD that are included as part of a major new release of the widely used *QSIPrep* software (Cieslak et al. 2021).

As described below, this update to *QSIPrep* provides major new preprocessing and post-processing features for HBCD. Major improvements to preprocessing address challenges in infant brain extraction, distortion correction, and normalization to infant-specific templates. Furthermore, we dramatically expanded the functionality of post-processing workflows as part of a new, dedicated software package – *QSIRecon. QSIRecon* produces rich derived data including diverse maps of tissue microstructure as well as person-specific white matter bundles. Using 529 dMRI sessions across two timepoints of the HBCD 1.0 release, we provide empirical support for the utility of the new preprocessing methods, describe variation related to data quality and scanner vendor, and demonstrate expected development of microstructure and white matter bundles. The public release of both this software and the landmark HBCD data will accelerate rigorous and reproducible white matter research on early life brain development.

## METHODS

For HBCD, the dMRI working group extended the *QSIPrep* software package, culminating in the 1.0.0 release. Like the other modality-specific pipelines utilized by HBCD, *QSIPrep* is a BIDS App (Gorgolewski et al., 2017) that is rigorously tested via continuous integration, documented and containerized, and adheres to the NMIND software quality standard (Kiar et al., 2023) at the gold level. Even before development for HBCD, *QSIPrep* has been widely-used (>39.5k downloads / 600k runs at the time of writing) with a completely open development cycle and active community support. As summarized in **Figure 1**, below we detail the specific improvements for HBCD. These include major improvements in preprocessing (as part of *QSIPrep*) as well as a completely new software package for post-processing reconstruction workflows (in *QSIRecon,* **Figure 2**).

**Figure 1.**
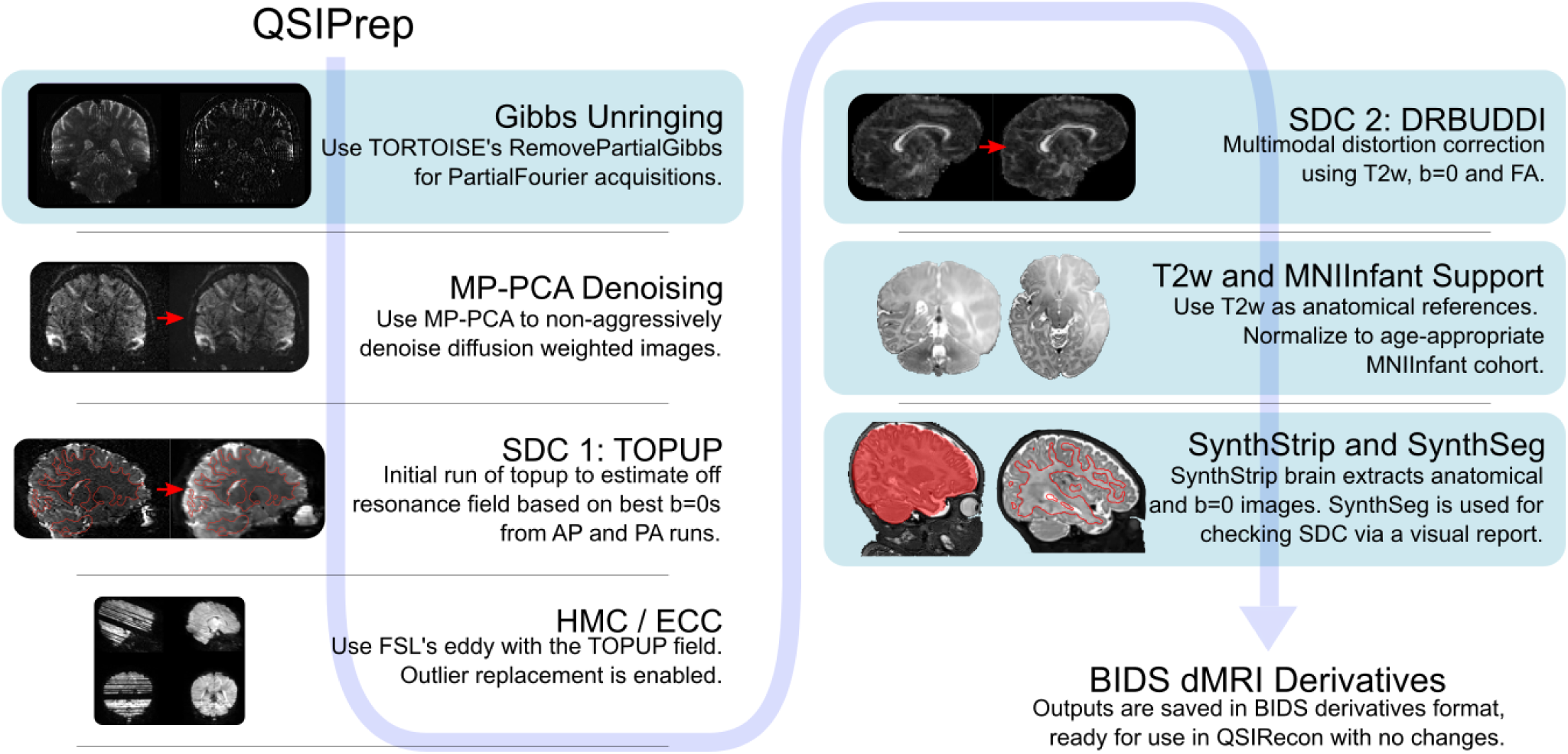
A diagram of the QSIPrep preprocessing workflow highlighting the features added for HBCD. Light blue shaded boxes indicate new features. The arrow indicates the order in which the steps are applied. The first two steps, the (newly added) partial Fourier Gibbs unringing is run first, followed by MP-PCA denoising from *MRtrix3*. An initial off resonance field is estimated from *b*=0 images from the AP and PA phase-encoding polarity scans by *TOPUP* and passed to eddy where head motion correction (HMC), eddy current correction (ECC), susceptibility distortion correction (SDC) and outlier replacement is applied. Next, the newly-added *DRBUDDI* is run on the eddy outputs for a multimodal second distortion correction step. A final interpolation is performed to apply the *DRBUDDI* correction and transform the dMRI into AC-PC space aligned with the T2w image. The transformation is computed between the AC-PC T2w image and a newly-supported age-appropriate MNIInfant template. The T2w image is brain extracted by the newly-added *SynthStrip*. Processed data are saved in BIDS derivative format.

**Figure 2.**
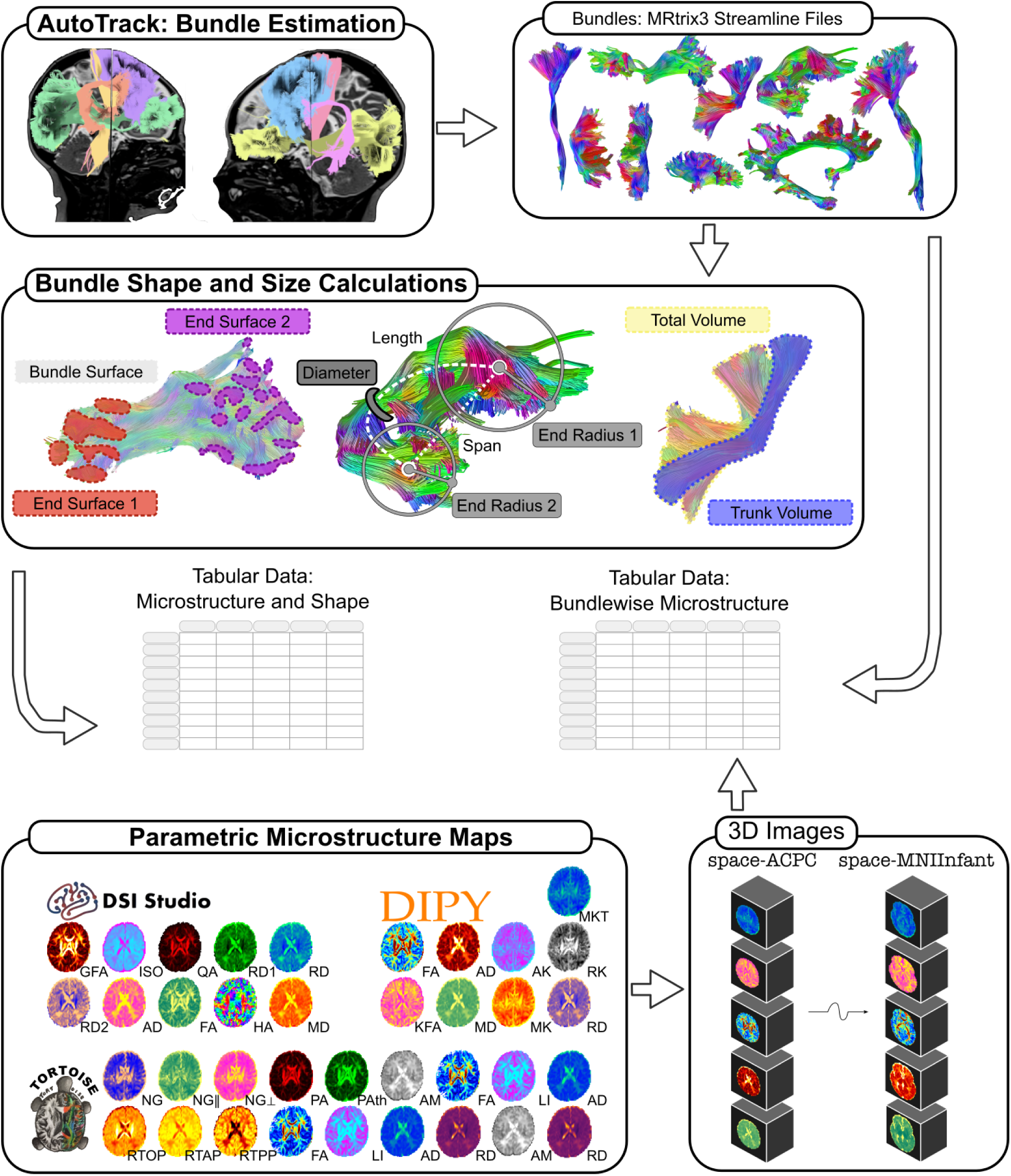
QSIRecon, a new BIDS app for postprocessing dMRI, was created for HBCD. Taking preprocessed dMRI BIDS derivatives as input, QSIRecon allows users to fit models to dMRI data, create parametric microstructure maps, perform tractography/tractometry and produce tabular data. Here the specific postprocessing workflow used for HBCD is shown. *DSI Studio*’s AutoTrack is first run to estimate a set of person-specific bundles. Bundle size and shape measures are calculated. Models from *DSI Studio* (GQI, inner shell tensor), *DIPY* (DKI) and *TORTOISE* (MAPMRI, inner and full shell tensors) are fit and parametric microstructure maps are created from each model. Output data are provided to users in the form of tabular data files (microstructure map statistics within each bundle, bundle shape/size measures), a *tck* format streamline file for each bundle and 3D images of each parametric microstructure map in both subject-native AC-PC space and in MNIInfant 1.7mm space.

### QSIPrep

Similar to the other infant-oriented preprocessing pipelines developed for the Baby Connectome Project (BCP) (Howell et al., 2019) and the Developing Human Connectome Project (dHCP) (Bastiani et al., 2019), *QSIPrep* uses recent versions of *FSL*’s *TOPUP* (Andersson et al., 2003) and *eddy* for head motion correction, susceptibility distortion correction (SDC), eddy current distortion correction, outlier replacement (Andersson et al., 2016) and movement-by-susceptibility correction (Andersson et al., 2017). However, HBCD pilot testing showed that *QSIPrep* needed more robust methods for infant dMRI in specific areas: brain extraction, SDC, spatial normalization, and partial Fourier Gibbs ringing correction. A detailed description of the preprocessing is included in **Supplementary Methods**.

#### Brain extraction and segmentation

Prior to HBCD*, QSIPrep* used *antsBrainExtraction* for skull stripping – a method based on registering a T1-weighted brain scan to the adult OASIS template (Marcus et al., 2007). HBCD scanning sessions in infants prioritize the collection of a T2-weighted (T2w) image, and do not always include a T1-weighted (T1w) image, so instead of attempting to register an infant brain to a much larger brain with a different contrast, we changed *QSIPrep’s* brain extraction method to *SynthStrip* (Hoopes et al., 2022). *SynthStrip* does not require spatial normalization and performs well on a wide range of input types. Furthermore, in pilot testing, we noticed that the 3-tissue segmentation (white matter, gray matter and CSF) from *FSL*’s *FAST* were not accurate enough on infant T2w scans, so we similarly replaced *FAST* with *SynthSeg* (Billot et al., 2023) from *FreeSurfer* 7.3.1. The visual reports produced by *QSIPrep* use these tissue segmentations to highlight SDC performance.

#### Susceptibility distortion correction

Following the approach of the Human Connectome Project, HBCD uses monopolar diffusion-weighting gradients for its dMRI acquisition. Monopolar sequences improve SNR (through lower achievable TE) at the cost of increased distortion. The standard in the neuroimaging community for SDC is *FSL*’s *TOPUP*, which was also the pre-HBCD default method in *QSIPrep*. However, *TOPUP* only corrects distortions apparent in the *b* = 0 images. This leads to undercorrection of high-anisotropy brain structures that only appear in the *b*> 0 images (Irfanoglu et al. 2015). To address this issue, the HBCD dMRI preprocessing also includes the *DRBUDDI* distortion correction method (Irfanoglu et al., 2015), which performs a multimodal distortion correction that makes use of Fractional Anisotropy (FA) maps, synthetic *b*= 0 images, and the high-resolution T2-weighted structural image. *DRBUDDI* is included in the *TORTOISE* software package (Irfanoglu et al., 2025). Notably, we extended *QSIPrep*’s functionality to enable *TOPUP, DRBUDDI* or a sequence of *TOPUP-eddy-DRBUDDI* to be used for SDC. In the visual reports produced by *QSIPrep*, users can see images before and after SDC; in these reports, the gray/white matter boundary from *SynthSeg’s* tissue segmentation is shown as a contour line (**Supplementary Figure 1**).

#### Infant-specific templates

It is critically important to choose an appropriate brain template to define a standard space for cross-participant image analysis (Avants et al., 2010). An ideal brain template represents the shape and appearance midpoint of the brains that will be registered to it. Too large of a template brain introduces an unintended smoothness to the resampled data, as single voxels in native space are spread across multiple voxels in the template space. Similarly, too small of a template introduces unintentional partial voluming effects when multiple native space voxels are potentially combined into single template space voxels. Prior to HBCD, the only supported template in *QSIPrep* was the MNI152NLin2009cAsym template, which is not appropriate for the age ranges scanned in ses-V02 (∼4.6 weeks postnatal, 40-44 weeks post-conceptional) and ses-V03 (∼25.2 weeks postnatal, 60-64 weeks post-conceptional). To address this, we added a feature to *QSIPrep* where the age of the participant at the time of scanning is read from the BIDS input and the appropriate MNIInfant template (Fonov et al., 2011) is selected as the target for spatial normalization based on the age at scan time. The MNIInfant template set includes cohorts for specific age ranges that have a more appropriate brain size, anatomy, and tissue contrast for each cohort. Derivative parametric maps produced by *QSIRecon* (see below) are then resampled into this automatically selected MNIInfant template.

While the age-specific atlases are beneficial for registration purposes, they make it more challenging to compare across ages. For the evaluations in this manuscript, we adopt the approach used by HBCD’s BOLD pipeline, NiBabies (Goncalves et al., 2025). Namely, the native to MNIInfant cohort nonlinear transform computed in *QSIPrep* via ANTs is combined with TemplateFlow’s template-to-template transform from the MNIInfant cohort to MNI152NLin6Asym space (Ciric et al. 2022).

#### MP-PCA denoising

Diffusion MRI is affected by random thermal noise during acquisition, affecting estimates of tissue microstructure and fiber orientation. Denoising can improve head motion correction performance (Cieslak et. al, 2024) and increases test-retest reliability of common dMRI derivatives (Manzano-Patron et al., 2023) at minimal loss of spatial resolution. Many modern dMRI preprocessing pipelines include image denoising as their first step (Theaud et al., 2020; Cai et al., 2021; Irfanoglu et al., 2025), including *QSIPrep*. Marchenko-Pastur principal components analysis denoising (MP-PCA; Veraart et al. 2016) is run on the AP and PA HBCD acquisitions separately using a 3-voxel window size. This is noteworthy in the context of HBCD because it is the first infant imaging consortium pipeline to include denoising (Bastiani et al., 2019; Howell et al., 2020).

#### Partial Fourier Gibbs ringing correction

Gibbs ringing is a common artifact in MRI that was previously addressed in *QSIPrep* by the *mrdegibbs* (Kellner et al., 2016) program from *MRtrix3* (Tournier et al., 2019). Although commonly used for all dMRI data, the algorithm in *mrdegibbs* was designed and validated specifically on data without partial Fourier acquisitions. HBCD’s dMRI acquisition has between 63-75% partial Fourier in the phase encoding direction, which has been shown to be better corrected by the “Remove Partial Gibbs” algorithm (Lee et al., 2021). We added the `--unringing-method rpg` option to use this method as implemented in *TORTOISE* (Irfanoglu et al., 2025).

### QSIRecon

As part of the changes introduced for HBCD, the reconstruction (postprocessing) workflows formerly included in *QSIPrep* were split into a separate software package – *QSIRecon* (https://doi.org/10.5281/zenodo.15013050). As part of developing the new *QSIRecon*, we created the `hbcd_scalar_maps` workflow to produce the derivatives distributed in the HBCD data release. This postprocessing workflow consists of two main parts – the calculation of whole-brain parametric microstructure maps and the estimation of a set of white matter bundles.

#### Whole-brain parametric microstructure maps

Voxelwise models fits of dMRI data can be used to produce spatial maps where each voxel’s value reflects a specific property of the diffusion process. The multi-shell dMRI acquisition of HBCD (with *b*-values of 0, 500, 1000, 2000, 3000 s/mm^2^, with *n* = 20, 12, 24, 36 and 60 volumes respectively) lends itself to a wide range of such dMRI models, which have different strengths and challenges in measuring specific aspects of the diffusion process. In order to accelerate both methodological and translational research, the HBCD workflow in QSIRecon includes four major models of the diffusion process. As each model often estimates several diffusion properties, at present these models yield over 40 whole-brain parametric microstructure maps per dMRI imaging session. A complete list of maps is presented in **Supplementary Table 2**. It should be noted that for the HBCD data release 1.0 we did not include models that assume fixed diffusivities – such as Restriction Spectrum Imaging (RSI; White et al., 2013) and Neurite Orientation Dispersion and Density Imaging (NODDI; Zhang et al., 2012) – as these fixed diffusivity values have been validated for adults, but vary dramatically in infancy and childhood (Guerrero et al., 2019). Below we describe the four models that are fit as part of HBCD’s postprocessing workflow.

#### Diffusion tensor imaging (DTI)

The diffusion tensor model (Basser et al., 1994a) – a fixture in dMRI – provides a simple way to describe Gaussian diffusion in a voxel. The tensor model has been used extensively in the human brain (Pierpaoli et al., 1996) including developmental neuroscience, with large known effects of increasing FA and decreasing MD with age (Qui et al., 2015). The eigenvectors and eigenvalues from the fitted tensor are used to calculate widely used scalar maps such as fractional anisotropy (FA), mean diffusivity (MD), axial diffusivity (AD) and radial diffusivity (RD). When fitting tensors, we adopt the approach from the Adolescent Brain and Cognitive Development (ABCD) study (Hagler et al., 2019) and perform multiple separate fits. One fit used only the low-*b* (*b* < 1000) inner shells, where the assumptions of the tensor model are most valid (de Santis et al. 2011) and the results will be more similar to legacy single shell studies. The second fit used all available data (e.g., *“full shell”*) as in Pines et al. (2020). The inner shell tensor fit is computed twice: once in DSI Studio using ordinary least squares and again in TORTOISE using weighted linear least squares (Basser et al., 1994b), where the tensor parameter estimation is weighted proportional to estimated SNR. The full shell fit is only done in TORTOISE with weighted linear least squares. Comparisons between full and inner shells should be done using the maps estimated by TORTOISE, while comparisons of tensor fitting methods can be done with DSI Studio and TORTOISE inner shell fits.

#### Diffusional kurtosis imaging (DKI)

Water diffusion in the brain is affected by the physical structures that make up neurons and organelles. Instead of freely diffusing through space, water encounters barriers from myelin, cell membranes and other structures that introduce non-Gaussian features into the water diffusion distribution. The Diffusional Kurtosis Imaging (DKI, Jensen et al., 2005) model extends the of the DTI model by adding an additional 15 parameters that capture the deviations from Gaussianity missed when fitting the simple 6 parameter DTI model. The DKI model incorporates data from all shells, potentially estimating the same scalar maps from DTI (FA, MD, etc) more accurately than a traditional tensor fit (Henriques et al., 2021). In addition to the measures from DTI, the DKI model also allows one to compute additional scalars derived from the kurtosis tensor such as mean kurtosis (MK), radial kurtosis (RK), and axial kurtosis (AK) (Jensen & Helpern, 2010). DKI’s sensitivity to non-Gaussian diffusion makes it useful for capturing the interaction of water with more complex tissue features.

#### Mean Apparent Propagator MRI (MAPMRI)

The Mean Apparent Propagator (MAPMRI) method (Özarslan et al., 2013) is a model-free approach that captures complex water diffusion. As a matter of practice, a diffusion tensor is first computed (using just the inner shells (*b*<1250), saved as an output) to determine the coordinate framework in which the ensemble average diffusion propagator (EAP) is to be estimated in three dimensions by Hermite polynomials. MAPMRI is estimated in *TORTOISE* (Irfanoglu et al., 2023) and maps are derived for multiple EAP-related properties. One set of maps captures the probability of a water molecule returning to its origin (RTOP) (which is inversely proportional to the pore size), to its principal axis (RTAP), or the plane perpendicular to the principal axis (RTPP) (which is inversely proportional to the analog of radial diffusivity). Furthermore, non-Gaussianity (NG) is calculated for the entire 3D space, along the principal direction of diffusion (NGPar) and perpendicular to it (NGPer). The anisotropy of the EAP is calculated as “propagator anisotropy” (PA). We also calculate the propagator anisotropy 1 (PAth), as the angular distance between the fitted MAPMRI coefficients and the coefficients corresponding to its isotropic version (Özarslan et al., 2013). Prior work in adolescents and young adults has shown that MAPMRI scalars are robust to head motion and among the most sensitive to age effects (Pines et al., 2020). Critically, our estimation of MAPMRI uses the metadata present in BIDS to define the Large and Small delta diffusion-gradient timing parameters (Δ and δ) for each scan^1^.

#### Generalized q-Sampling Imaging (GQI)

A key feature of the post-processing in the HBCD pipeline is the use of generalized *q*-sampling imaging (GQI). GQI is another model-free approach that estimates water diffusion orientation distribution functions (dODFs) using an analytic transform of the diffusion signal (Yeh et al., 2010). Like the other methods, GQI produces a number of parametric microstructure maps such as generalized fractional anisotropy (GFA), quantitative anisotropy (QA), and isotropic component (ISO) (Yeh et al., 2013).

We chose to use the peak directions of dODFs estimated via GQI as the basis for tractography instead of the popular constrained spherical deconvolution (CSD) method for a number of practical reasons. Although the HBCD dMRI acquisition meets the criteria to use the multi-shell multi-tissue (MSMT) CSD method, the age range sampled by HBCD makes it difficult to apply the method consistently and optimally. For instance, MSMT-CSD works optimally on two tissue types in neonates (Pietsch et al., 2019; Grotheer et al., 2022), but typically three tissues are included when analyzing images from adults (Jeurissen et al., 2014). Also, GQI does not require a response function and is applied identically regardless of age. Finally, while GQI has been used extensively with adult data over the last 15 years, it has also been used successfully for tractography in infants (Borchers et al., 2020; Dennis et al., 2019; J. Y. Lee et al., 2021; Barnes-Davis et al., 2024) including data from dHCP (Sun et al., 2024).

#### Estimation of person-specific white matter bundles

The AutoTrack implementation in *DSI Studio* (version “Chen”) was used to estimate a set of canonical bundles from the preprocessed dMRI. We selected AutoTrack as it performed well on pilot data and has recently been used with dMRI data from infants in the same age range as ses-V03 (∼25.2 weeks) in HBCD (Zhang et al., 2025). It should be emphasized that many other high-quality bundle extraction methods exist. However, many current methods have technical limitations related to the age range and acquisition protocol collected by HBCD. For example, methods like Automated Fiber Quantification (AFQ; Yeatman et al. 2012; Kruper et al., 2021) depend on waypoint and exclusion masks in a template space. There currently is no set of T1w or T2w templates with waypoint/exclusion masks appropriate for the full age range sampled by HBCD. Furthermore, issues (described above) about different optimal ODF methods would likely require different methods at different timepoints. In contrast, probabilistic methods like XTRACT (Warrington et al., 2020) do not produce streamlines, which are necessary to produce shape and complexity metrics available for streamline bundles. Furthermore, distance bias in probabilistic tractography may also interact with the dramatic changes in brain size. Finally, many methods – including both AFQ and RecoBundles (Garyfallidis et al., 2018) – require a complete tractogram as input, which would require a single set of step size and turning angles be picked *a priori*; this would be difficult to choose for the age range of HBCD.

AutoTrack begins with a set of rigorously-defined anatomical bundles, defined here as sets of streamlines following the spatial trajectory of an established projection in the brain, in a template space. Three dimensional QA and ISO images in the same template space are also provided. A GQI fit is performed on subject native preprocessed dMRI data from which QA+ISO maps are estimated. A multimodal nonlinear registration is computed between the participant QA+ISO maps and the corresponding templates. The template space bundle streamlines are then warped into subject native space via these transforms. Person-specific bundles are estimated by generating streamlines in subject native space over a range of reasonable deterministic tracking parameters (including step size and turning angle); they are assigned to a bundle if their geometry is within a specified Hausdorff distance of a transformed template bundle. Shape, size, and geometry measures (Yeh, 2020) are calculated for each bundle and distributed in tabular format. General size measures include length, volume, and surface area, which are calculated along with measures at bundle endpoints such as endpoint surface area, radius, and irregularity. Shape measures of bundle curl (a measure of overall curvature) and elongation (a quantification of “straightness”) reflect the overall shape of each bundle. Concrete examples of each are shown in **Figure 2**. Furthermore, all (40+) parametric measures of microstructure described above are also averaged within each bundle (both weighted and unweighted by streamline density), yielding a rich feature set. All bundle quantification is performed in subject native AC-PC space.

### Application to HBCD

Here we assess the HBCD dMRI pipeline using actual HBCD data. We ran *QSIPrep* version 1.0.0 and *QSIRecon* version 1.0.1 on the HBCD 1.0 release BIDS data. To ensure no major errors were introduced, we performed manual visual inspection of the raw and preprocessed data, followed by a quantitative comparison of automated QC scores before and after processing. As detailed below, we specifically assessed the effect of processing on measures of image quality, the impact of brain masking with *SynthStrip,* the effect of applying MP-PCA and Gibbs unringing, and the benefits of distortion correction with *DRBUDDI*. The parametric microstructure maps and bundles were also visually inspected. We ran voxelwise mass univariate analyses on the derived data to replicate known effects of brain development and describe scanner vendor differences in microstructure estimates. AutoTrack bundle sizes were compared between sessions along with bundle averages of microstructure estimates.

#### Data

The HBCD beta release dataset (*n*=529 sessions) includes data from the first two sessions to include MRI scanning (V02 and V03). **Table 1** shows the ranges of ages per scanning session. All data was collected with parental consent (Dean et al., 2024). This cohort was primarily from Siemens scanners. **Supplementary table 1** details the number of scan sessions per HBCD site included in the present analysis, and lists the scanner model and manufacturer in use at each site.

**Table 1:**
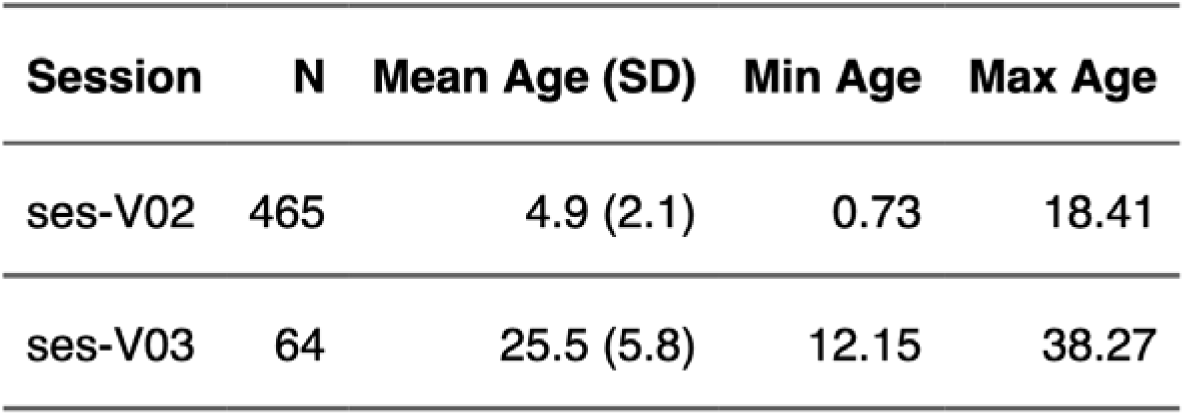
Sample statistics from the HBCD beta release dMRI data. Age is in weeks.

The HBCD dMRI acquisition is described in detail elsewhere (Dean et al., 2024). Briefly, it includes two complete multishell dMRI acquisitions with opposite phase encoding directions (AP and PA). The AP and PA acquisitions sample the same shells (*b*= 0,500,1000,2000,3000 s/mm^2^) with different subsets of diffusion-sensitizing directions, producing two unique, complete sampling schemes with *n* = 10, 6, 12, 18 and 30 volumes in each shell respectively (per acquisition). We chose to acquire paired scans with opposite phase encoding polarity with diffusion-weighting in each so that distortion correction could incorporate *b*> 0 data, which we ultimately accomplish using *DRBUDDI*. The dMRI scan sampled 87 1.7mm contiguous 140×140-voxel axial slices with an FOV of 238×238 mm^2^ at multiband factor 3 and no additional in-plane acceleration. The dMRI scans used the same TR (4800 ms), TE (88 ms), and similar partial Fourier sampling in the phase encoding direction (63–75%) across vendors.

#### Evaluation of automated QC measures

Automated QC measures are useful for identifying serious issues with dMRI data. *QSIPrep* calculates a suite of 34 automated QC metrics. For this application to HBCD, we focused on three: contrast to noise ratio (CNR), the Neighboring DWI Correlation (NDC), and the coregistration Dice distance. CNR is calculated per shell by *eddy* and has been used as an outcome measure for comparing image denoising methods (Patron et al., 2024) and comparing acquisitions across scanner manufacturers (Harms et al., 2025). CNR is calculated in each voxel as the standard deviation of the Gaussian Process (GP) predictions from *eddy* divided by the standard deviation of the residuals (the difference between the observations and the GP predictions). A median CNR value is calculated per shell within the brain mask, which we used to compare CNR distributions across vendors and shells.

NDC is the mean spatial correlation between closest images in *q*-space. Prior work has demonstrated that NDC (Yeh et al., 2019) tracks closely with the quality ratings of dMRI experts (Richie-Halford et al. 2022); it has also been used as a proxy for dMRI data quality (Meisler et al., 2022; Verhelst et al., 2025; Sydnor et al. 2025). In our initial validation of *QSIPrep*, we showed that NDC improves with preprocessing (Cieslak et al., 2021) and with accurate head motion correction (Cieslak et al., 2024). Here we repeat this experiment with the updated pipeline as a vital validity check. All instances of NDC plotted/analyzed in this manuscript use the unmasked version of the calculation. Both masked and unmasked NDC is available in the tabular QC files included in *QSIPrep* outputs.

The coregistration Dice distance describes the overlap of the *SynthStrip* masks for the T2w brain and the *SynthStrip* mask derived from the mean *b*=0 image after coregistration. High values indicate low overlap between the two. We evaluated this QC measure prior to voxelwise analyses of scalar maps in template space.

#### Evaluation of SynthStrip

Brain masks calculated by *SynthStrip* on the T2w images during *QSIPrep* preprocessing were transformed into MNI152NLin6Asym space at 1.7mm isotropic resolution. Group coverage maps were calculated separately for ses-V02 and ses-V03 due to the large difference in brain size and contrast. We calculated coverage maps as the fraction of times a given voxel was included in the brain mask (transformed into atlas space via nearest neighbor interpolation), to highlight areas where there is disagreement among individual masks. In these maps, values close to .5 indicate high variability, while those close to 1 (or 0) indicate that *SynthStrip* consistently included (or excluded) that area from the brain mask across all participants.

#### Evaluating the effect of denoising and Gibbs unringing

As the first major consortium pipeline to include Gibbs unringing and MP-PCA, we evaluated the impact of these steps on data quality. To this end, we ran *QSIPrep* without MP-PCA and Gibbs unringing while keeping all other settings identical. Specifically, we compared NDC and CNR, as both values are calculated after denoising and unringing are performed.

#### Evaluating susceptibility distortion correction

The largest change from the standard *QSIPrep* workflow is the use of *DRBUDDI* for SDC. In order to evaluate the effect of using *DRBUDDI*, we additionally preprocessed this cohort without *DRBUDDI* (but otherwise identically) and compared both FA and NDC. For the FA comparison, we adopt the approach used in Irfanoglu et al. (2015) in which the AP and PA data are split after preprocessing and then a DTI model is fit separately for each set of phase encoding directions. The FA from the AP and PA DTI models are then subtracted and the absolute difference is warped to MNINLin6Asym space for comparison. Misalignment between high-FA structures is highlighted in the group average FA absolute difference maps. We also compared NDC between the TOPUP+DRBUDDI and TOPUP-only runs. We did not evaluate CNR, as it is calculated prior to the application of DRBUDDI.

#### Bundle evaluation

AutoTrack produced streamline data in MRtrix3 `tck` format and tabular data of bundle shape statistics (Yeh, 2020). Bundles from a small random selection of subjects were plotted in 3D along with the brain mask to check for appropriate size, shape and location of bundles. Summaries of shape data were plotted across bundles and sessions to verify that bundle volume, region area, and irregularity change as expected between sessions V02 and V03. Scalar values within each bundle are also plotted between sessions.

#### Statistical analyses

Some parametric scalars, such as FA and MD, are known to show large developmental effects in the age range of the data considered here. As a validity check on the pipeline outputs, we used a simple linear model to estimate the effects of using age, vendor, and data quality (summarized by the raw Neighboring DWI Correlation) using the *ModelArray* package (Zhao et al., 2023). Mass univariate testing used scalar maps transformed to MNI152NLin6Asym space via a single interpolation by combining the MNIInfant warps from preprocessing with the appropriate template-to-template transform from *TemplateFlow*. It should be emphasized that technical studies from working groups (such as the HBCD dMRI working group) that use pre-release data are explicitly not allowed to test specific developmental hypotheses. In that context, all developmental effects are presented as a validity check; detailed nonlinear modeling (e.g., using generalized additive mixed models) for developmental inference are beyond the scope of the present methodological effort.

## RESULTS

### Qualitative evaluation

Many of the imaging artifacts affecting dMRI, including susceptibility-related distortion, movement-related slice drop out, and thermal noise, are clearly visible in raw dMRI images. Accordingly, we first visually inspected raw and corrected images to ensure the corrections are being applied appropriately. A clear example is displayed in **Figure 3**, where the processed images are undistorted, less noisy and less affected by slice drop-out. Due to the on-scanner B1-(receive-field) bias correction, there is no obvious intensity inhomogeneity in the images.

**Figure 3.**
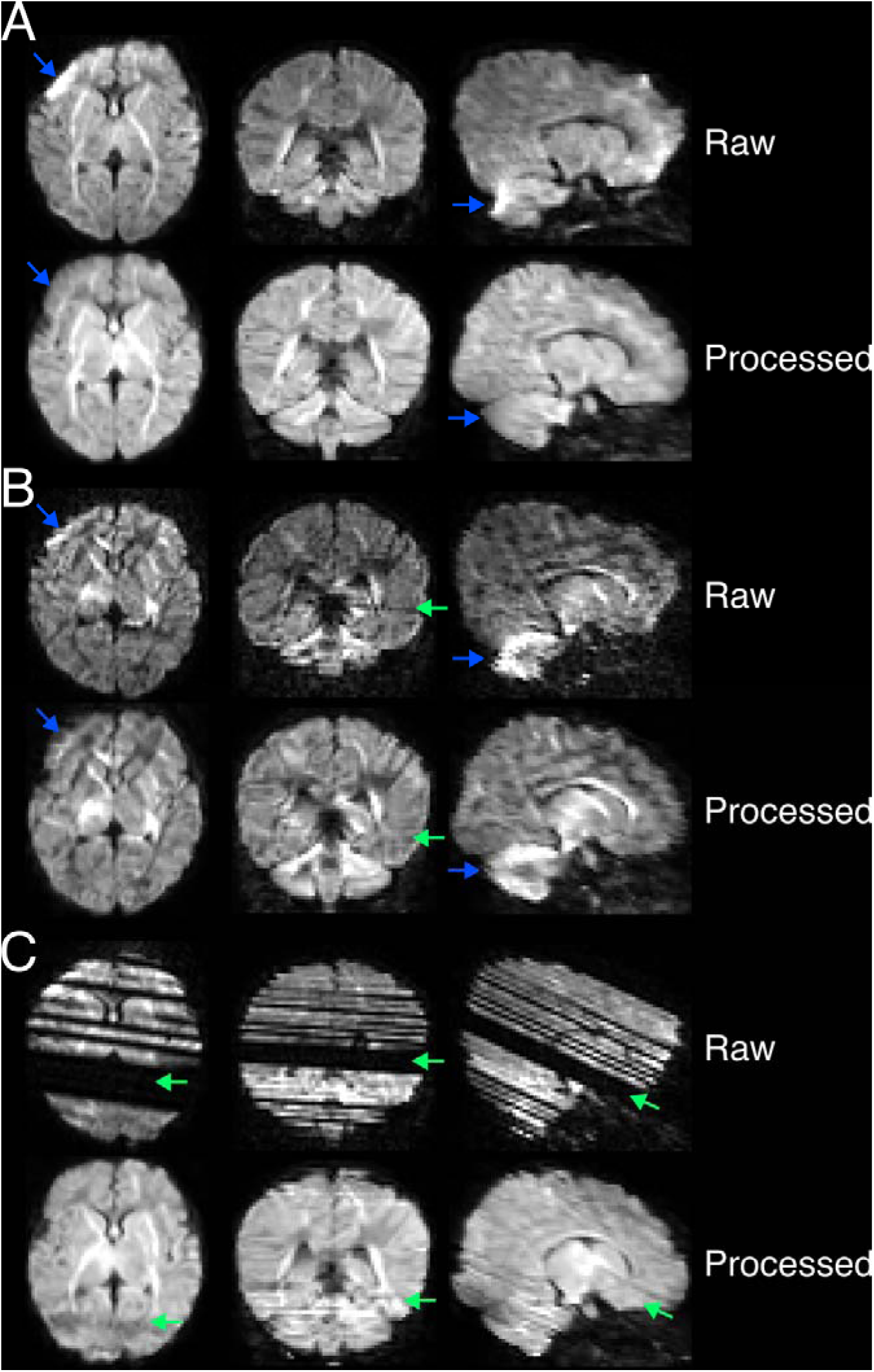
Examples of image artifacts before (top) and after (bottom) correction in 3 volumes collected on the Siemens Prisma platform running the Syngo MR XA30 operating system in the P-A phase encoding direction. To facilitate comparison, raw images were resampled into approximate rigid alignment with their preprocessed version, which is displayed in the bottom row of each panel. Note that due to this resampling, the distortion in the raw images is no longer purely in the P-A direction, resulting in the appearance of the raw data being spatially larger. Across all panels, susceptibility distortion is highlighted by blue arrows and motion-related slice dropout by green arrows. **A.** Slices from a *b*=1000 volume of a relatively artifact-free acquisition from the scan with P-A polarity only exhibit susceptibility distortion and its accompanying signal pileup in the frontal lobe, both corrected in the processed volume. **B.** Slices from a *b*=2000 volume in the same scan exhibit similar susceptibility distortion and pileup, but also some slice dropout. The outlier slice replacement in *eddy* appears to be quite accurate. **C.** Slices from a *b*=1000 volume in which the raw data was severely affected by slice dropout to such an extent that the distortion-related pileup is barely visible. The corrected images are dramatically improved (due to outlier replacement), but artifacts are still visible.

### Image processing improves measures of data quality

We began by examining the Contrast to Noise (CNR) values for each shell, separately for each scanner manufacturer (**Figure 4A**). Notably, none of the distributions are heavily skewed toward zero – suggesting that the gradient directions are being handled correctly and that the Gaussian Process modeling within *eddy* is successful at finding signal in each shell. CNR summary statistics are presented for each scanner model in **Supplementary Table 3**. Notably, Siemens scanners had highest CNR at all *b* values, followed by Phillips and then GE. Next, we evaluated how preprocessing changes an important measure of dMRI quality – the Neighboring DWI Correlation (NDC) – which summarizes how similar a given volume is to its neighbors in *q*-space. In general, higher quality images have higher NDC, and preprocessing should improve NDC. As expected, NDC increases following preprocessing (**Figure 4B**) for all scanner models. Together, these results suggest that the HBCD dMRI acquisition is sensitive to real dMRI signal and that the artifacts affecting dMRI signal are being mitigated during preprocessing.

**Figure 4.**
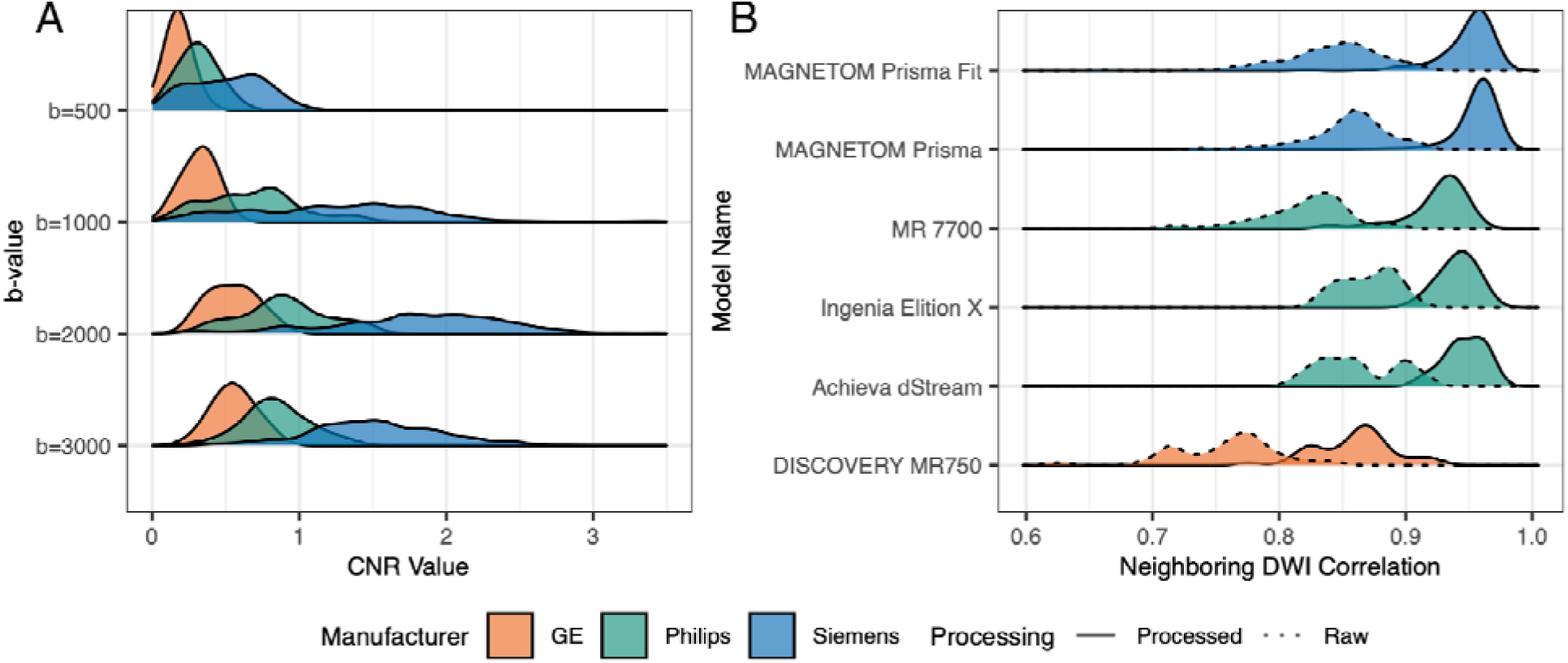
Automated measures of image quality: CNR and Neighboring DWI Correlation (NDC). **A.** Contrast to noise ratio (CNR) per shell and vendor is plotted. **B.** Kernel density plots are shown for unmasked NDC calculated before and after image processing, grouped by scanner manufacturer. Distributions of the raw data NDC are outlined by dashed lines and preprocessed by solid lines. NDC is increased after processing. Scanner model-specific CNR values are listed in **Supplementary Table 3**.

### SynthStrip robustly estimates brain masks in both sessions

The small size and rapidly changing tissue contrast of the infant brain makes brain extraction an ongoing challenge. Initial pilot testing showed that tools included in prior versions of QSIPrep frequently failed on infant brains. While *SynthStrip* was not specifically designed for infant brains, it has been shown to work effectively on many contrasts. As such, we evaluated *SynthStrip* in ses-V02 (age 0-1 months) and ses-V03 (age 3-6 months) data. Coverage maps of *SynthStrip* (**Supplementary Figure 2**) in both ses-V02 and ses-V03 shows that brains were accurately extracted by *SynthStrip* in the entire population. No area of the brain appears to be systematically missed and no areas outside the brain are consistently covered across the cohort. Notably, future releases of HBCD data will include masking performed by the Baby and Infant Brain Segmentation Neural Network (BIBSNET), which may further improve brain extraction (Hendrickson et al., 2025).

### Gibbs unringing and MP-PCA denoising increase QC scores

We evaluated the effect of MP-PCA and Gibbs unringing on CNR and NDC. As seen in **Figure 5**, including these two preprocessing steps markedly improved both NDC and CNR values. Inclusion of MP-PC and Gibbs unringing increased NDC values. Furthermore, CNR improvements were most apparent in shells with higher *b* values.

**Figure 5.**
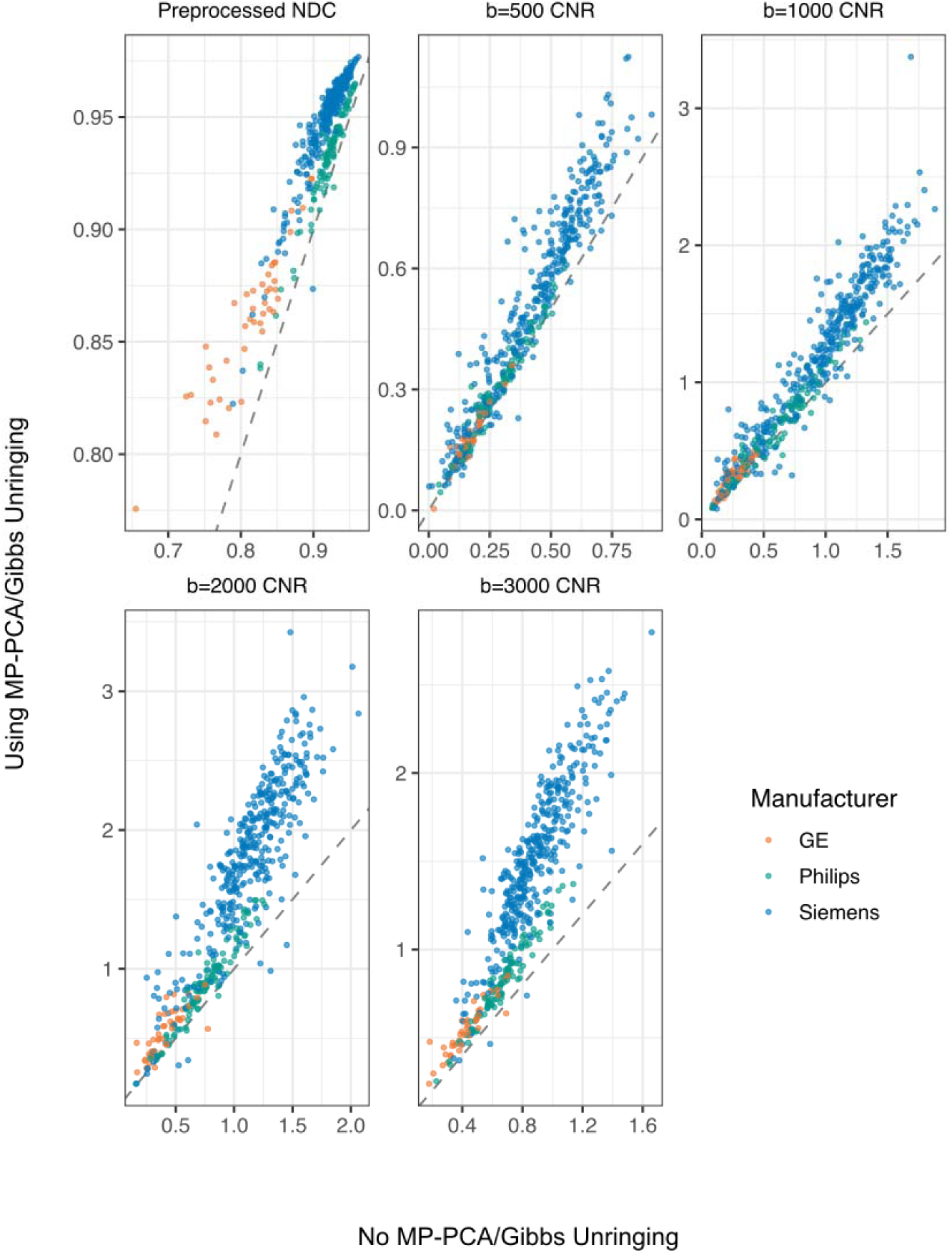
Contrast to Noise Ratio (CNR) and Neighboring DWI Correlation (NDC) increase when Gibbs unringing and MP-PCA denoising are included in preprocessing. The identity is plotted with a dashed line. The largest increases in CNR are seen for Siemens scans at.

### DRBUDDI improves distortion correction over TOPUP alone

Successful SDC should result in accurate spatial alignment of brain structures regardless of their phase encoding direction. Thus, a comparison of FA images computed separately from the AP vs. PA scans should show only minor, random differences. In **Figure 6** we see that using *TOPUP* without *DRBUDDI* results in clear misalignment between high-FA structures such as the internal capsule and the cerebellar peduncles. The difference is visually striking for both ses-V02 and ses-V03. This misalignment is reduced when *DRBUDDI* is used jointly with *TOPUP.* There is no statistical difference in NDC between the two versions (**Supplementary Figure 3**).

**Figure 6.**
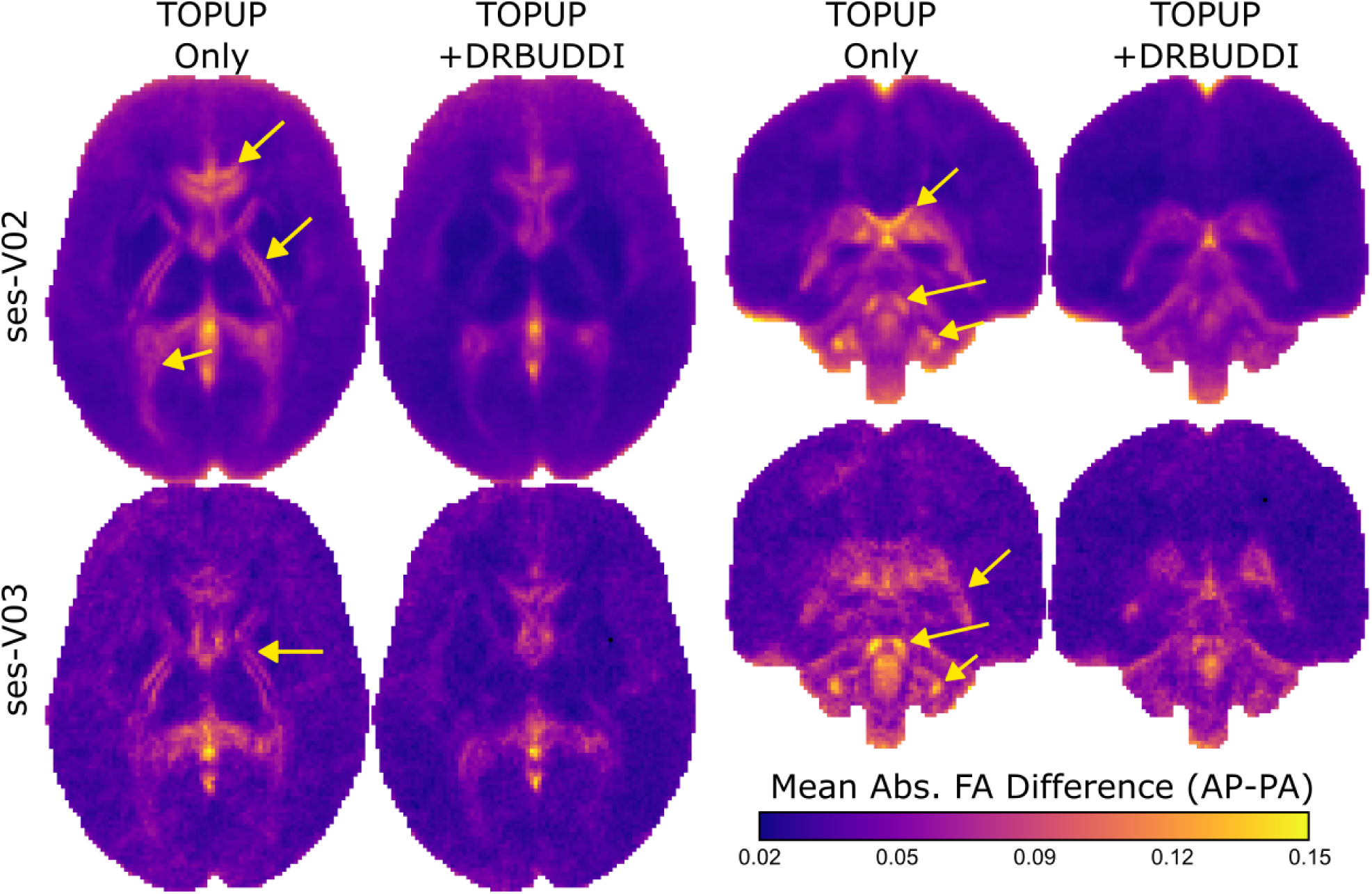
*TOPUP+DRBUDDI* better aligns high-FA structures in AP and PA scans compared to TOPUP alone. Mean absolute difference in FA calculated from only AP and only PA scans is shown for both sessions V02 (average across n=465 sessions) and V03 (average across n=64 sessions). Misalignment in AP and PA scans of high-FA structures creates bright spots in this map. *TOPUP*-only distortion correction reveals misalignment of the internal capsule and the optic radiation in the axial slices and the cerebellar peduncles in the coronal slices (indicated by arrows), which is notably reduced, although not entirely eliminated, by the inclusion of *DRBUDDI*.

### Parametric microstructure maps show expected associations with age, vendor differences and data quality

While detailed analyses of infant brain development are beyond the scope of this software paper, replication of well-established associations between age and derived measures like FA or MD provides a valuable validity check – evidence that preprocessing, modelling and spatial normalization have all functioned properly. To this end, we evaluated associations with age on parametric microstructure maps using mass univariate linear models (**Figure 7, Supplementary Figure 4**). We first evaluated registration quality as quantified using the coregistration Dice distance. This revealed that a subset of 67 sessions from ses-V02 had inadequate coregistration (see **Supplementary Figure 5**), most commonly due to the infant not being scanned in a supine position. We removed these 67 sessions from the mass univariate statistical analyses; results including these participants are displayed in **Supplementary Figure 6**.

**Figure 7.**
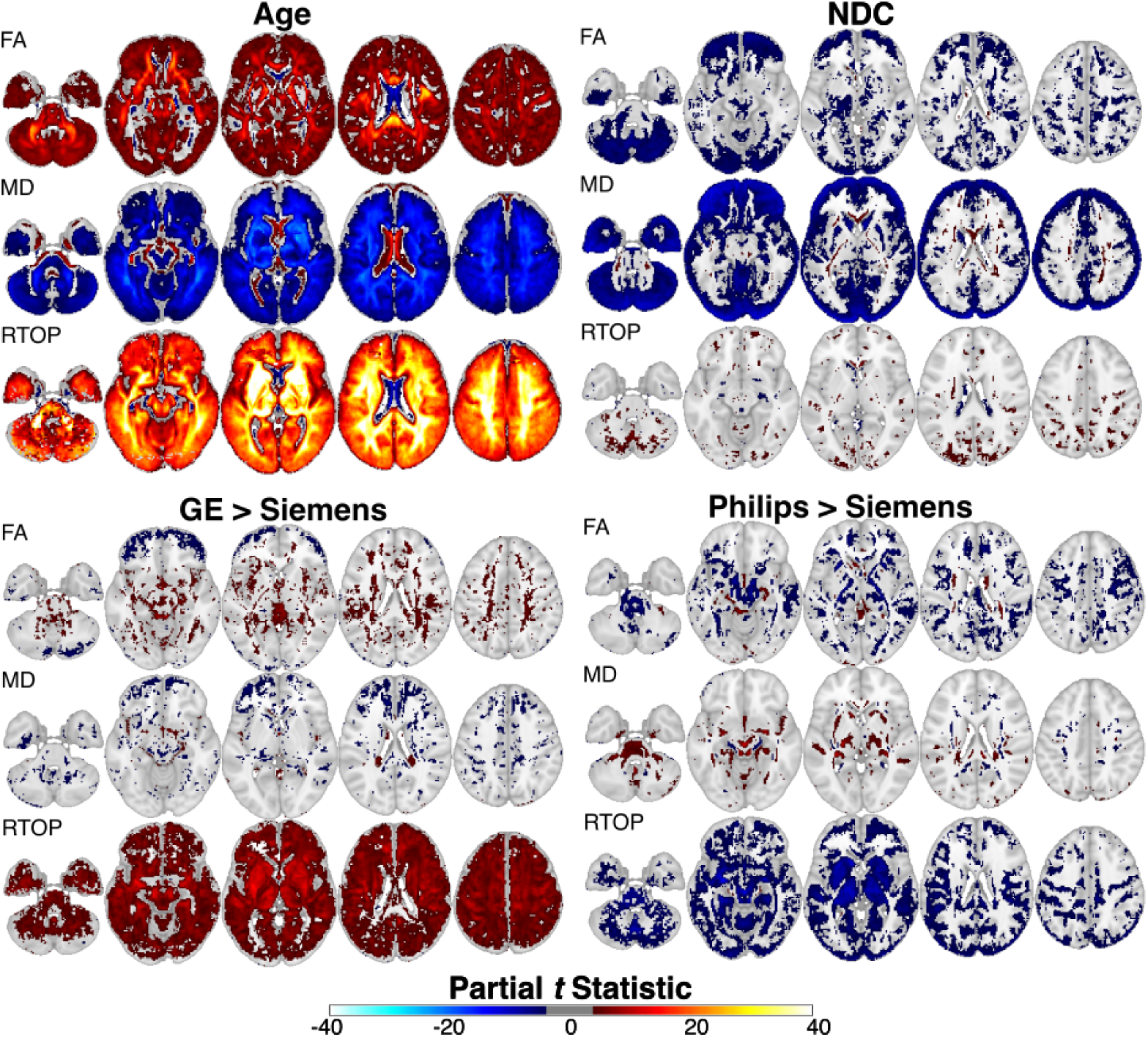
Linear models were fit for FA, MD and RTOP, where each was modeled as a function of age, NDC and scanner manufacturer. Sessions included had a coregistration Dice distance less than 0.06 (ses-V02 *n*=399, ses-V03 *n*=63). Partial t statistics, plotted as color overlays on the MNI152NLin6 template brain, assess the significance of individual predictor variables by testing the effect of one predictor while controlling for all other predictors in the model. Age and NDC are continuous predictors, with one coefficient (and one partial *t*) estimated for each testing against 0. FA and RTOP increase with age, while MD decreases – a replication of findings from other studies. NDC *t* maps show that FA and MD are affected by data quality in large spatially contiguous regions while RTOP is significantly affected only in smaller regions. Manufacturer is a categorical variable where the partial *t* tests for significance compared to the mean value for Siemens. RTOP is most affected by manufacturer difference, with GE estimating higher RTOP and Philips estimating lower RTOP across most of the brain. An exploratory threshold of -3<*t* <3 are left transparent.

As reported in prior studies, we found that FA increases during the first postnatal year of life, while MD declines (Qui et al., 2015; Ouyang et al., 2019;). Notably, we also found that a measure of water restriction from the MAPMRI model – the return to origin probability (RTOP) – also increases with age, with effect sizes that are appreciably higher than are seen for traditional tensor-based measures like FA or MD.

Our linear model also allowed us to simultaneously quantify the effects of data quality and manufacturer differences while accounting for age. As above, we summarized data quality using NDC, where lower values indicate lower quality. We found in general that lower NDC – likely related to participant motion (see **Supplementary Figure 7**) – was associated with higher FA near the edge of the brain. In contrast, RTOP was less affected by variation in data quality. While these analyses suggest that RTOP may be more sensitive to age and less sensitive to data quality than traditional measures like FA and MD, we also found that RTOP had more widespread associations with variation in scanner vendor.

### AutoTrack yields person-specific white matter bundles

Generating high quality tractography requires accurate preprocessing, including correct gradient rotation, brain masking, and dODF estimation. A qualitative examination of identified bundles shows that the AutoTrack workflow in *QSIRecon* appropriately estimates dODFs (**Supplementary Figure 8**) and captures bundle shape and location (**Figure 8**). Within individuals, the bundles show consistent shape and location between the two scanning sessions. The size and complexity of the bundle terminations appears to increase in the second session.

**Figure 8.**
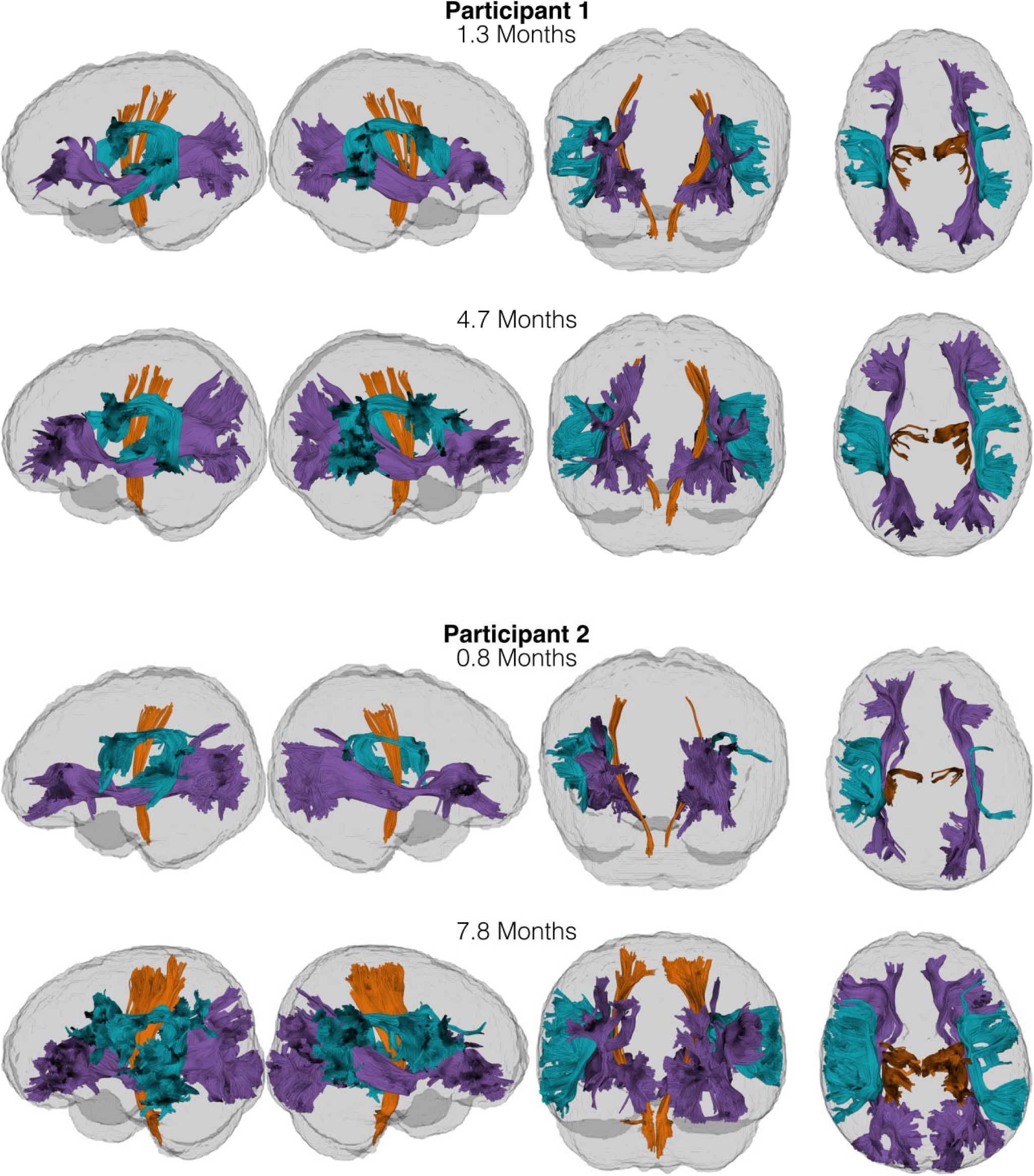
Participant-specific major white matter bundles are plotted for two longitudinal participants. The arcuate fasciculus (turquoise), corticospinal (orange) and the inferior frontal occipital fasciculus (purple) are seen in both sessions of both participants. Views of the left hemisphere and right hemisphere bundles are followed by posterior and superior views of bundles from both hemispheres. The gray surface contains the session’s SynthStrip mask.

While **Figure 8** shows data from two representative participants, **Figure 9** summarizes bundle shape data from all the sessions in the beta release. Notably, observations from the two example participants generalize to the rest of the sample: bundle volume, termination surface area and shape irregularity increase from ses-V02 to ses-V03.

**Figure 9:**
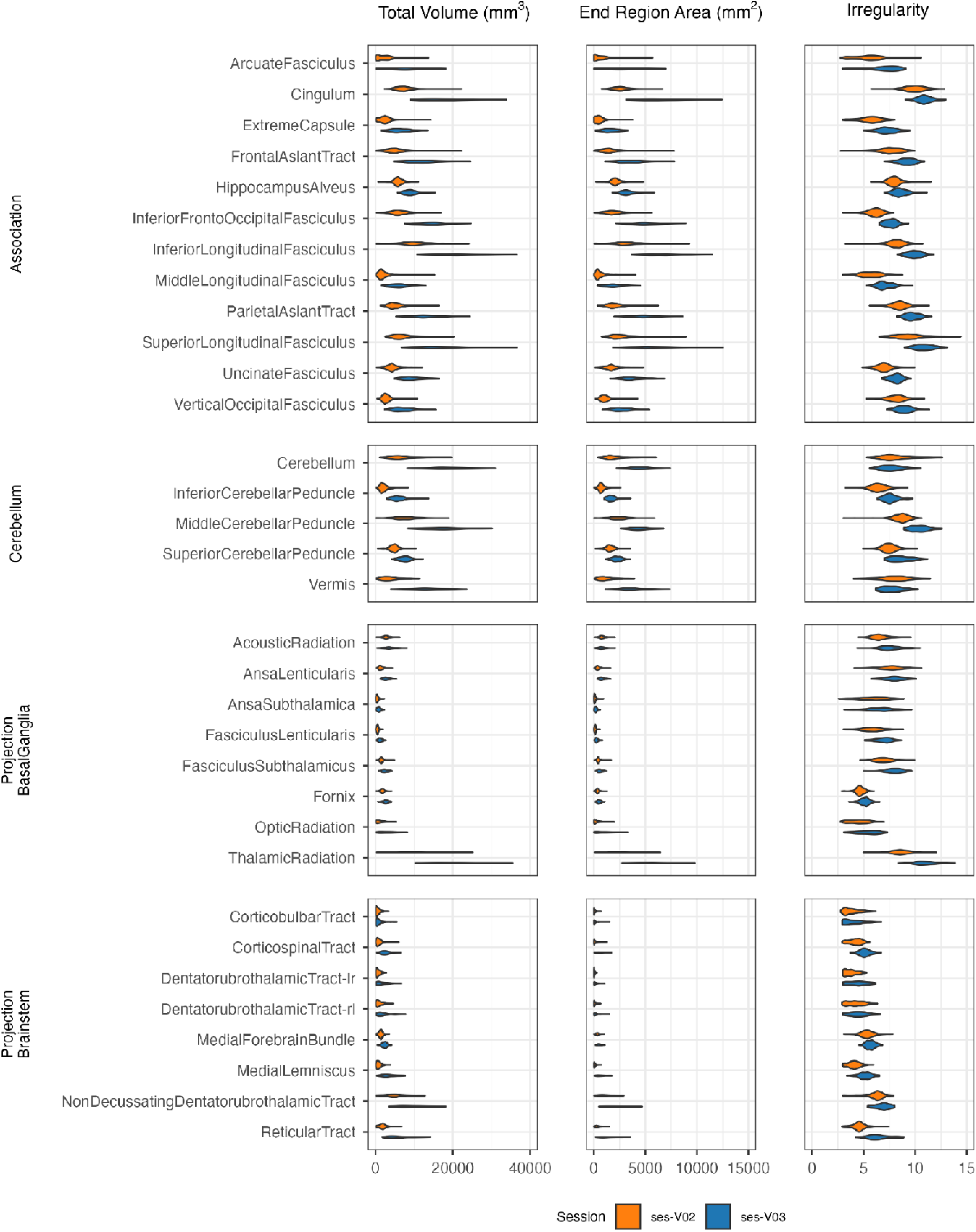
Bundles increase in volume, expand their cortical termination area and become more irregular over the 2-6 months between ses-V02 and ses-V03. These changes are expected with brain growth and white matter development. Violin plots for each bundle, collapsed across hemispheres, are shown for both sessions. The corpus callosum bundles, corticostriatal and corticopontine bundles were omitted from the plot because they are significantly larger than the other bundles. The -rl and -lr suffixes denote the hemispheres where the Dentatorubrothalamic bundle begins and ends, respectively.

Recall that bundles can also be used similarly to how parcels are used in cortical segmentations – as masks over which to aggregate parametric microstructure maps. Therefore, we expected to similarly see age-related changes in FA and MD across sessions in the bundle microstructure means. **Figure 10** shows all the bundles and three of the many scalars that can be summarized within them. As expected, the same developmental patterns seen in the voxelwise analysis are clearly apparent: mean FA in the bundles increases between sessions, while MD declines. As in the voxelwise analysis, we observed even greater changes in RTOP between sessions than those seen in FA or MD.

**Figure 10.**
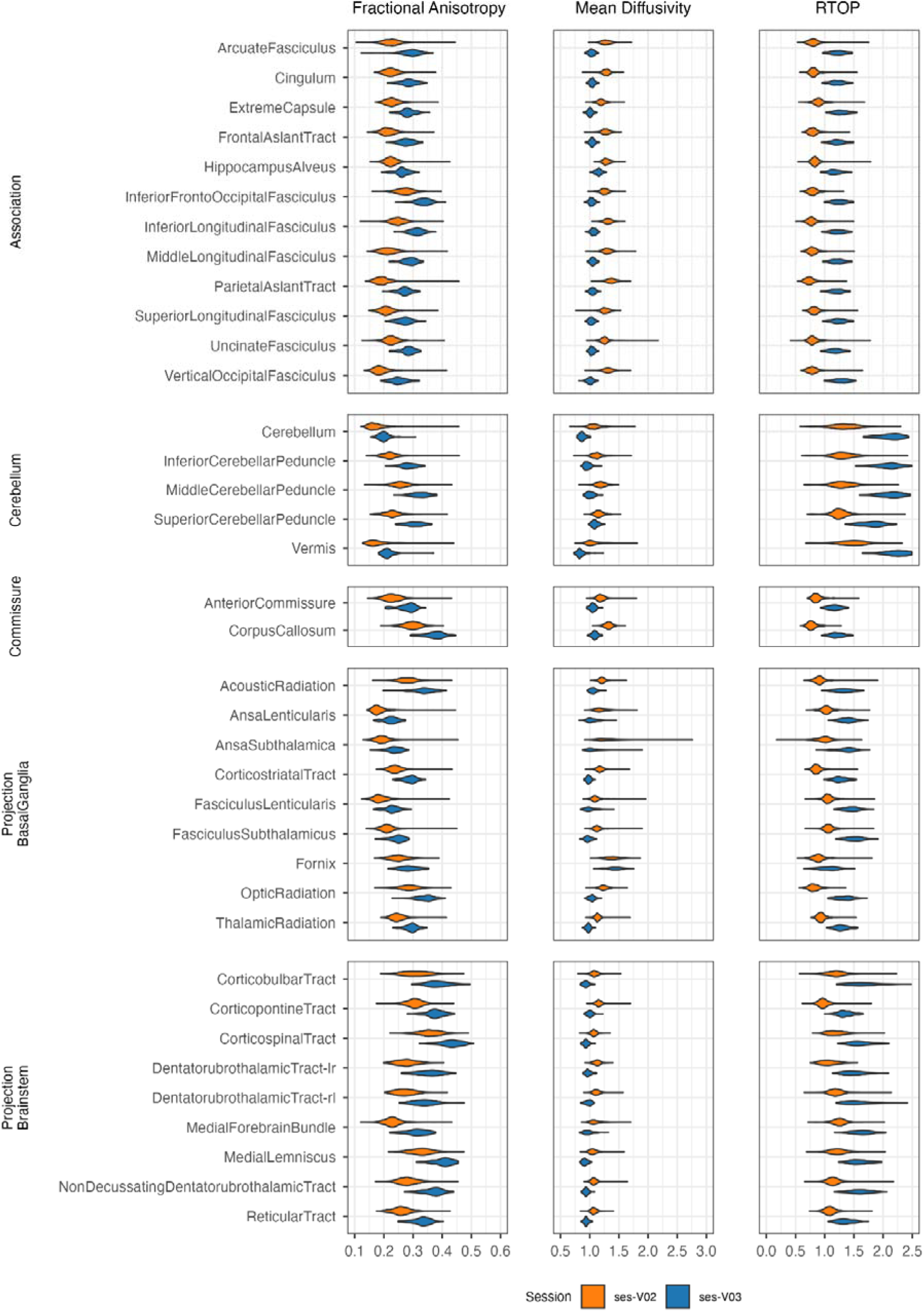
FA and MD show the characteristic patterns of increasing and decreasing with age, respectively. RTOP shows a similar effect to FA, but with a much higher effect size. Violin plots for each bundle, collapsed across hemispheres, are shown for both sessions.

## DISCUSSION

The HBCD project will scan individuals across a period of life when the brain is undergoing massive, rapid developmental change. To measure these changes in white matter more accurately, we built new features into *QSIPrep*, introduced the *QSIRecon* package, and applied them to data from HBCD.

Similar to dMRI pipelines from many other consortia (Glasser et al. 2013; Bastiani et al., 2019, Howell et al., 2019), *QSIPrep* uses *TOPUP* and *eddy* for SDC, head motion correction, eddy current correction and outlier replacement. However, it also adds MP-PCA denoising, *DRBUDDI* distortion correction, partial Fourier Gibbs unringing and MNIInfant registration targets. These additional steps respectively enable better dMRI-to-anatomical correspondence, higher fidelity signal around tissue compartment boundaries, and promote higher quality mass-univariate analyses in a standard-space. Manual qualitative visual inspection and comparison of quantitative automated QC scores showed expected improvements with preprocessing. Notably, the within-brain CNR estimated by *eddy* shows vendor differences similar to those reported in other multi-vendor consortia (Harms et al., 2025), which will need to be considered in downstream analyses (e.g., Figure 7). Quantitative comparison showed that inclusion of MP-PCA and Gibbs unringing in the pipeline improved both CNR and NDC. Similarly, quantitative comparison of processing with and without the second SDC step provided by *DRBUDDI* revealed that high FA structures from the AP and PA scans are better aligned when *DRBUDDI* is applied. This extends the benchmarking results in Irfanoglu et al. (2015) to infants.

To accommodate the interaction of postprocessing features desired for HBCD, we introduced *QSIRecon.* While postprocessing BIDS apps have existed for fMRI for years (Mehta et al., 2024), *QSIRecon* is to our knowledge the first to gather multiple software packages that are already widely used in the community and apply them with image acquisition metadata to preprocessed dMRI BIDS derivatives. For HBCD, the reconstruction workflow is quite inclusive – yielding over 40 parametric maps – but does not include all possible derived measures supported by *QSIRecon^2^*. Most notably, we do not include pairwise regional connectivity measures, as these remain controversial due to systematic false positives (Maier-Hein et al., 2017). For most researchers using data from HBCD, the derivative data from *QSIRecon* most appropriate for their study question can be downloaded directly from https://nbdc-datashare.lassoinformatics.com – saving computation time and storage burden. Researchers who want to explore models not included in the HBCD *QSIRecon* workflow can download the minimally preprocessed data (from *QSIPrep*) and fit any model they prefer.

Replication of firmly-established associations between age and both FA and MD provides evidence of the quality of the preprocessing workflow and derived measures generated by *QSIRecon.* Additionally, as previously observed in adolescents (Pines et al., 2020), we found that advanced measures like RTOP from MAPMRI are both more sensitive to brain development and robust to variations in data quality. These results underscore the importance of contemporary multi-shell dMRI acquisitions and recent advances in dMRI signal modeling for studies of brain development in early life.

There are limitations to the work presented here. First, more accurate coregistration is needed for some sessions in ses-V02. In the meantime, users can use the available coregistration Dice distances to exclude sessions with underestimated brain masks. Second, we used a simple linear age model to estimate age effects. While the replication of basic prior findings regarding age is a useful validity check, detailed developmental modeling is beyond the scope of this paper and will no doubt be a major focus of researchers who access the public data. Third, we also note that a successful run through *QSIPrep* and *QSIRecon* does not guarantee that the data are of adequate quality; focused quality control efforts are ongoing and remain necessary. Fourth, uniform preprocessing by *QSIPrep* and *QSIRecon* does not obviate the need for statistical harmonization. Vendor and site differences will require careful application of harmonization tools (Beer et al., 2020; Chen et al., 2022) and/or be considered as part of the statistical modeling. Finally, the slice-to-volume feature in ‘eddy’ for estimation and correction of intra-volume motion is not enabled for HBCD due to unavailable slice timing information on the Philips platform.

Moving forward, we anticipate that this software and the data generated as part of HBCD will accelerate studies of brain development in early life and beyond. Notably, other than MNIInfant support, all the features added to *QSIPrep* and *QSIRecon* are also applicable to adult dMRI data and are likely to prove beneficial for dMRI across the lifespan. Furthermore, the set of scalars calculated for HBCD will also be provided as part of the forthcoming release of the ABCD BIDS Community Collection (Feczko et al. 2021) version 7 (Meisler et al., in prep), creating a consistent parameter space for studying white matter development from infancy to adulthood.

## Data and code availability

QSIPrep is available at https://hub.docker.com/r/pennlinc/qsiprep, with source code at https://github.com/PennLINC/qsiprep. QSIRecon is available at https://hub.docker.com/r/pennlinc/qsirecon, with source code at https://github.com/PennLINC/qsirecon. Code written for this paper is available at https://github.com/PennLINC/hbcd_pipeline_replication_guide. The HBCD data used for this paper can be accessed via https://www.nbdc-datahub.org.

## Conflicts of interest

Andrew L. Alexander: Editor - Imaging Neuroscience, Scientific Advisor - ImgGyd, LLC Damien A. Fair is a patent holder on the Framewise Integrated Real-Time Motion Monitoring (FIRMM) software. He is also a co-founder of Turing Medical Inc that licenses this software. The nature of this financial interest and the design of the study have been reviewed by two committees at the University of Minnesota. They have put in place a plan to help ensure that this research study is not affected by the financial interest. This interest has been reviewed and managed by the University of Minnesota in accordance with its Conflict of Interest policies.

## Declaration of generative AI and AI-assisted technologies in the writing process

Generative AI was not used in any form in the writing of this manuscript.

## Footnotes

1 Since common DICOM to NIFTI conversion tools do not currently include diffusion timing parameters in the sidecar json files, the values were manually added.

2 A complete list of available workflows and their derivatives can be found at https://qsirecon.readthedocs.io/en/latest/builtin_workflows.html

## SUPPLEMENTARY MATERIAL

**Supplementary Table 1:**
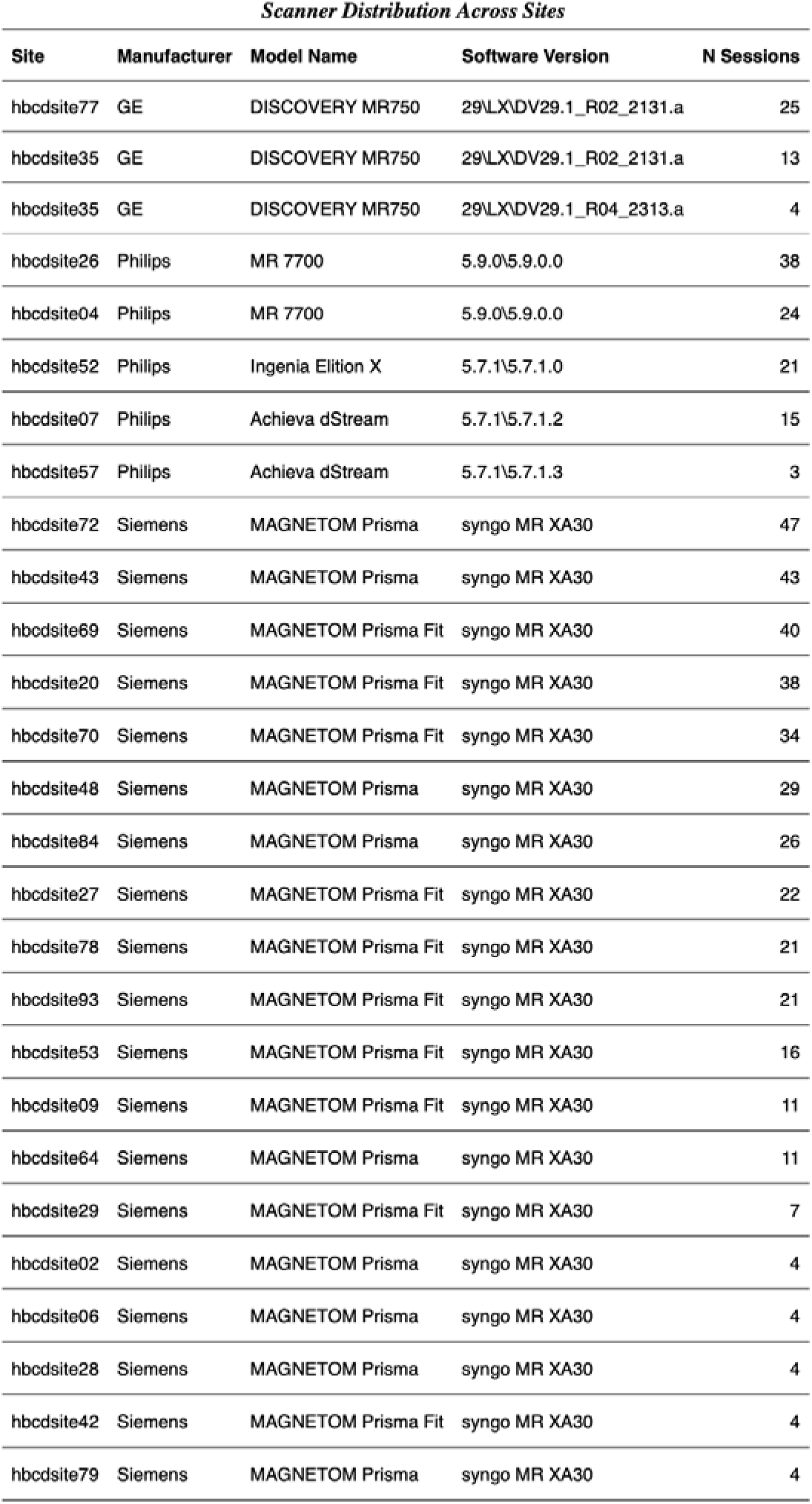
Sources of the sessions from the HBCD beta release. If a site upgraded their scanner OS or model, it appears in multiple rows (e.g., site35).

**Supplementary Table 2.**
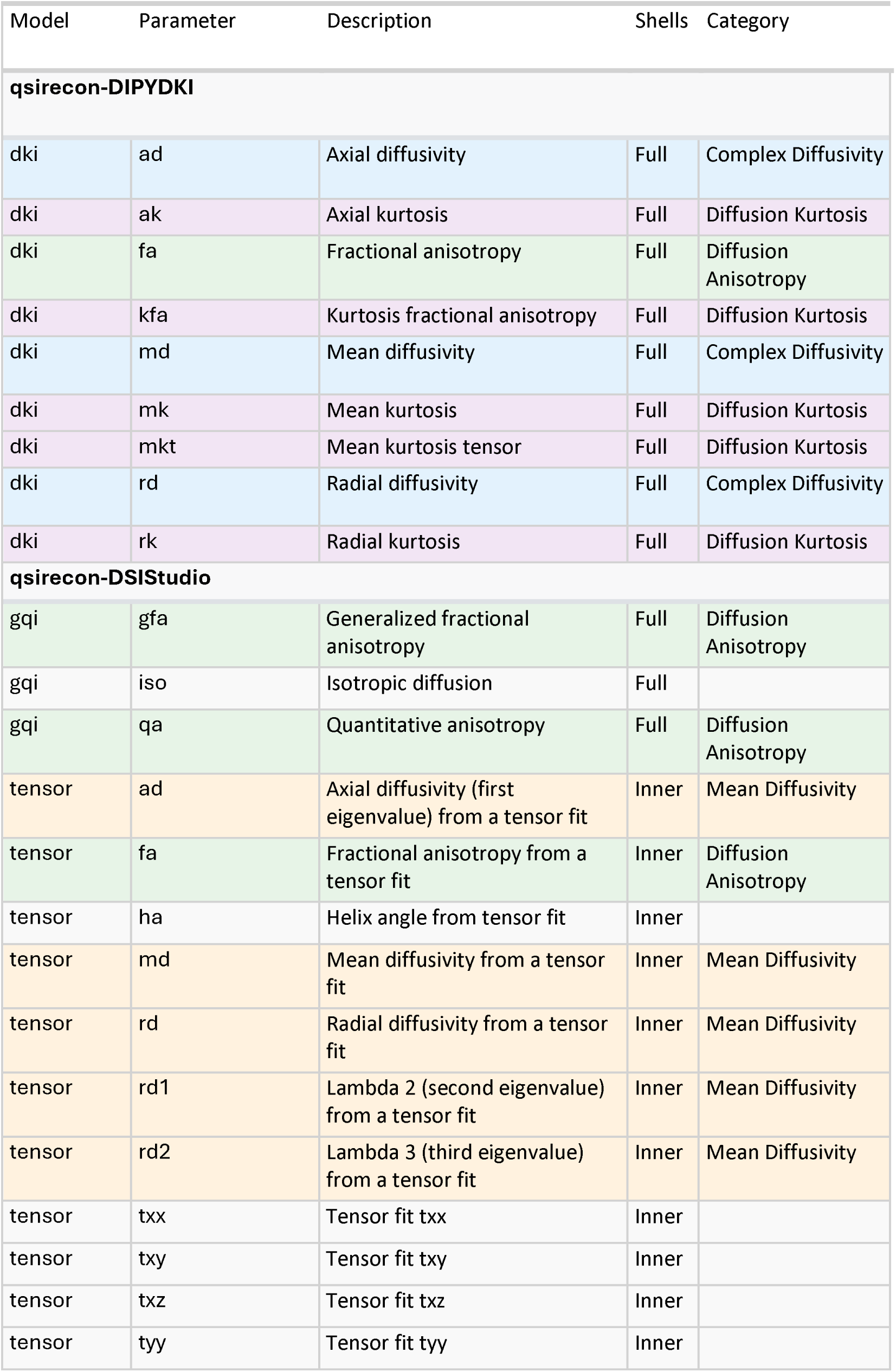
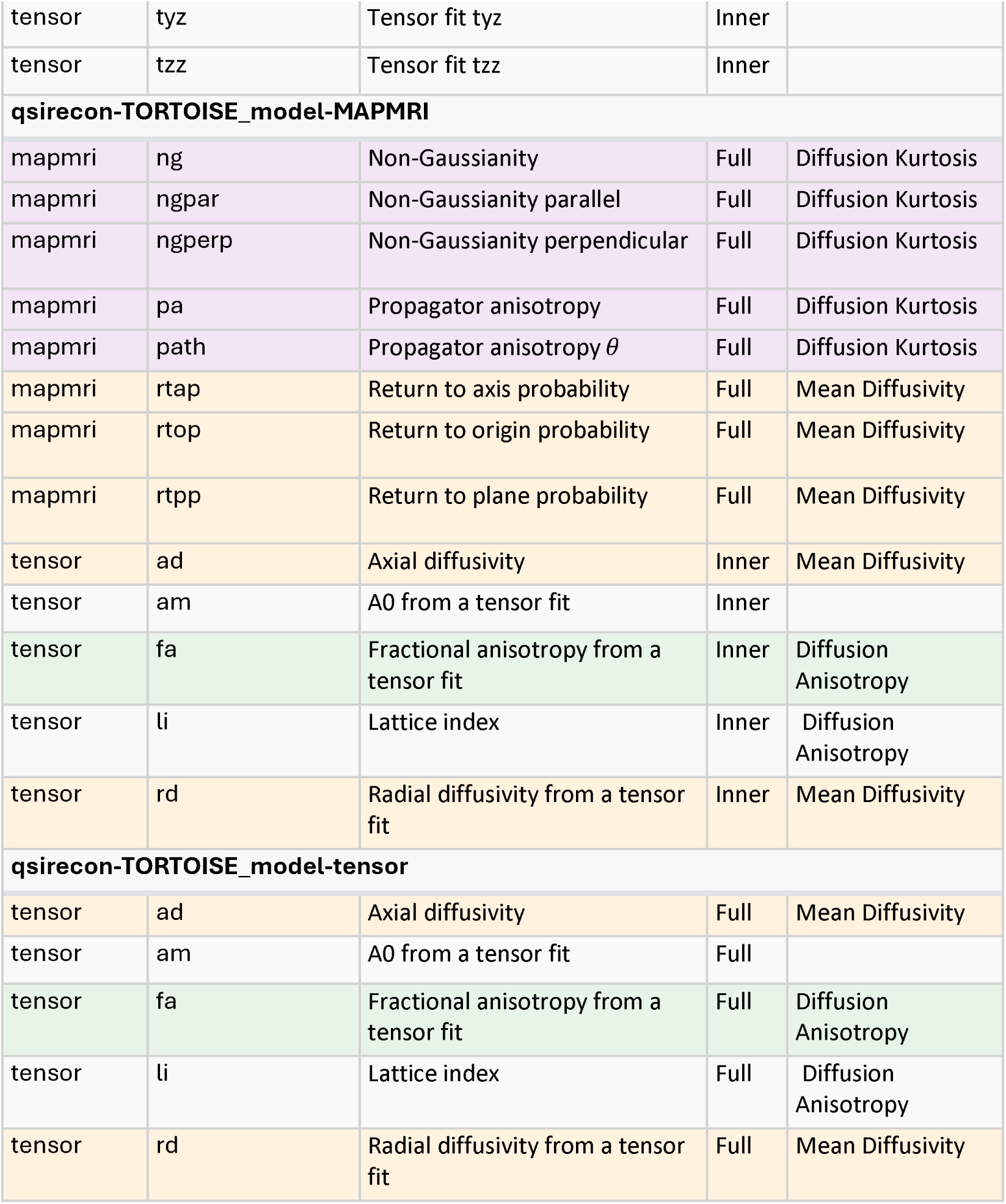
Parametric microstructure maps produced by *QSIRecon* from the hbcd_scalar_maps workflow. Where applicable, the categorization from Sadikov et al., (2025) is included as a guideline for general interpretation of each parameter (each Category is shaded with a different color).

**Supplementary Table 3.** Model-specific CNR values for each shell.

| Manufacturer | Manufacturer Model Name | b-value | Mean CNR | Median CNR | Std CNR |
| --- | --- | --- | --- | --- | --- |
| GE | DISCOVERY MR750 | b=3000 | 0.556 | 0.555 | 0.142 |
| Philips | Achieva dStream | b=3000 | 1.036 | 1.044 | 0.203 |
| Philips | Ingenia Elition X | b=3000 | 1.000 | 1.007 | 0.179 |
| Philips | MR 7700 | b=3000 | 0.741 | 0.747 | 0.183 |
| Siemens | MAGNETOM Prisma | b=3000 | 1.630 | 1.611 | 0.625 |
| Siemens | MAGNETOM Prisma Fit | b=3000 | 1.453 | 1.434 | 0.416 |
| GE | DISCOVERY MR750 | b=2000 | 0.544 | 0.548 | 0.165 |
| Philips | Achieva dStream | b=2000 | 1.071 | 1.147 | 0.310 |
| Philips | Ingenia Elition X | b=2000 | 1.182 | 1.242 | 0.263 |
| Philips | MR 7700 | b=2000 | 0.761 | 0.837 | 0.222 |
| Siemens | MAGNETOM Prisma | b=2000 | 1.927 | 2.039 | 0.557 |
| Siemens | MAGNETOM Prisma Fit | b=2000 | 1.679 | 1.731 | 0.550 |
| GE | DISCOVERY MR750 | b=1000 | 0.320 | 0.340 | 0.103 |
| Philips | Achieva dStream | b=1000 | 0.662 | 0.562 | 0.345 |
| Philips | Ingenia Elition X | b=1000 | 0.932 | 0.873 | 0.364 |
| Philips | MR 7700 | b=1000 | 0.606 | 0.626 | 0.257 |
| Siemens | MAGNETOM Prisma | b=1000 | 1.316 | 1.406 | 0.529 |
| Siemens | MAGNETOM Prisma Fit | b=1000 | 1.157 | 1.183 | 0.551 |
| GE | DISCOVERY MR750 | b=500 | 0.176 | 0.171 | 0.063 |
| Philips | Achieva dStream | b=500 | 0.318 | 0.285 | 0.155 |
| Philips | Ingenia Elition X | b=500 | 0.395 | 0.402 | 0.138 |
| Philips | MR 7700 | b=500 | 0.288 | 0.318 | 0.102 |
| Siemens | MAGNETOM Prisma | b=500 | 0.561 | 0.604 | 0.239 |
| Siemens | MAGNETOM Prisma Fit | b=500 | 0.484 | 0.489 | 0.249 |

**Supplementary Figure 1.**
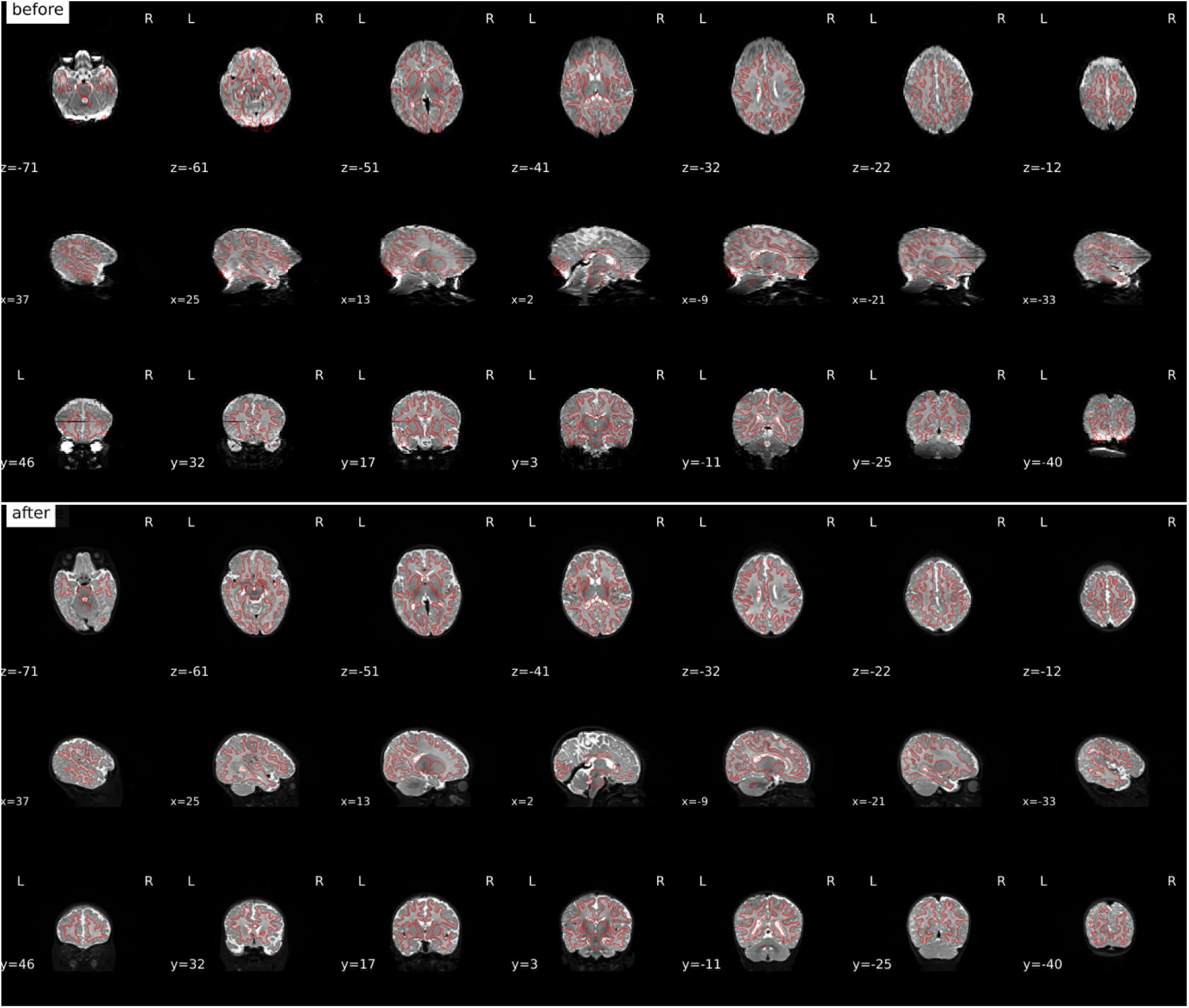
An example visual report of the *SynthSeg* white matter segmentation overlaid on a *b*=0 image before and after correction by *TOPUP*. While not used quantitatively in *QSIPrep* or *QSIRecon*, the segmentation is a useful guide to visually inspect the accuracy of SDC.

**Supplementary Figure 2.**
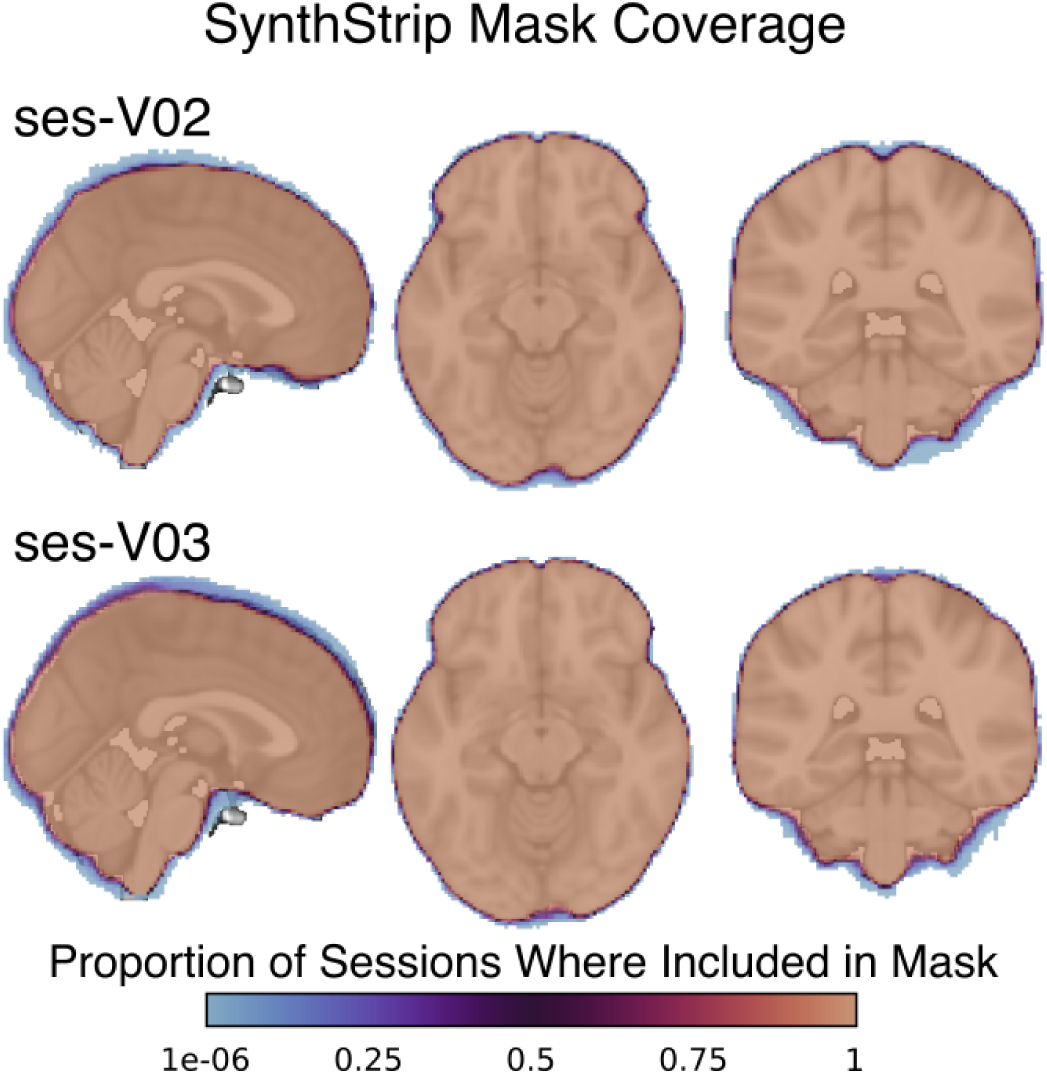
The coverage map of infant T2w-based *SynthStrip* masks mapped to MNINLin6Asym space. The highest variability of whether a voxel is masked or not is in a tight ribbon around the cortical surface. Both sessions V02 and V03 show high agreement on which voxels are within the brain.

**Supplementary Figure 3.**
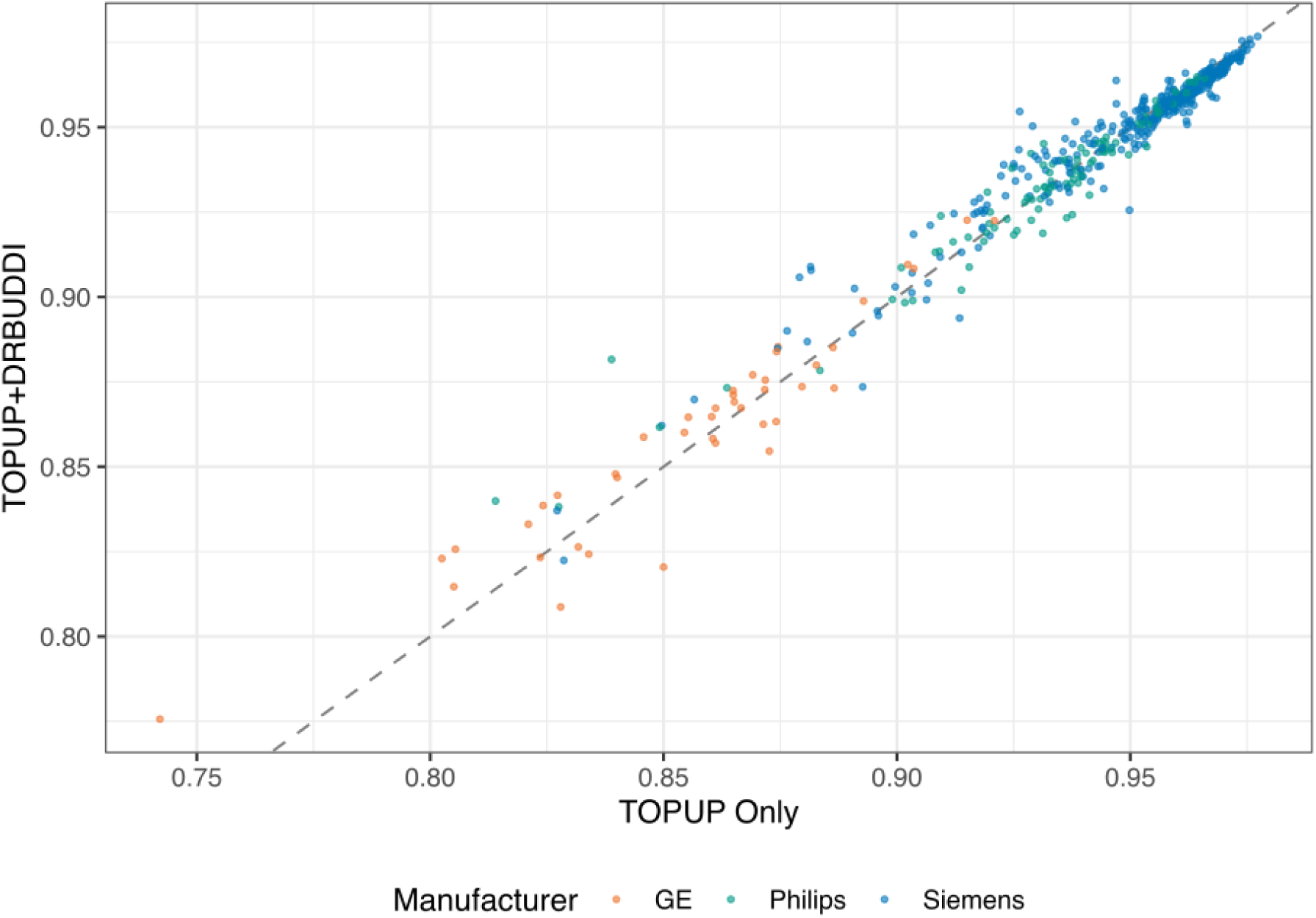
There is no significant difference in NDC when including DRBUDDI in preprocessing paired *t*(528) = -0.94 *n.s*.

**Supplementary Figure 4.**
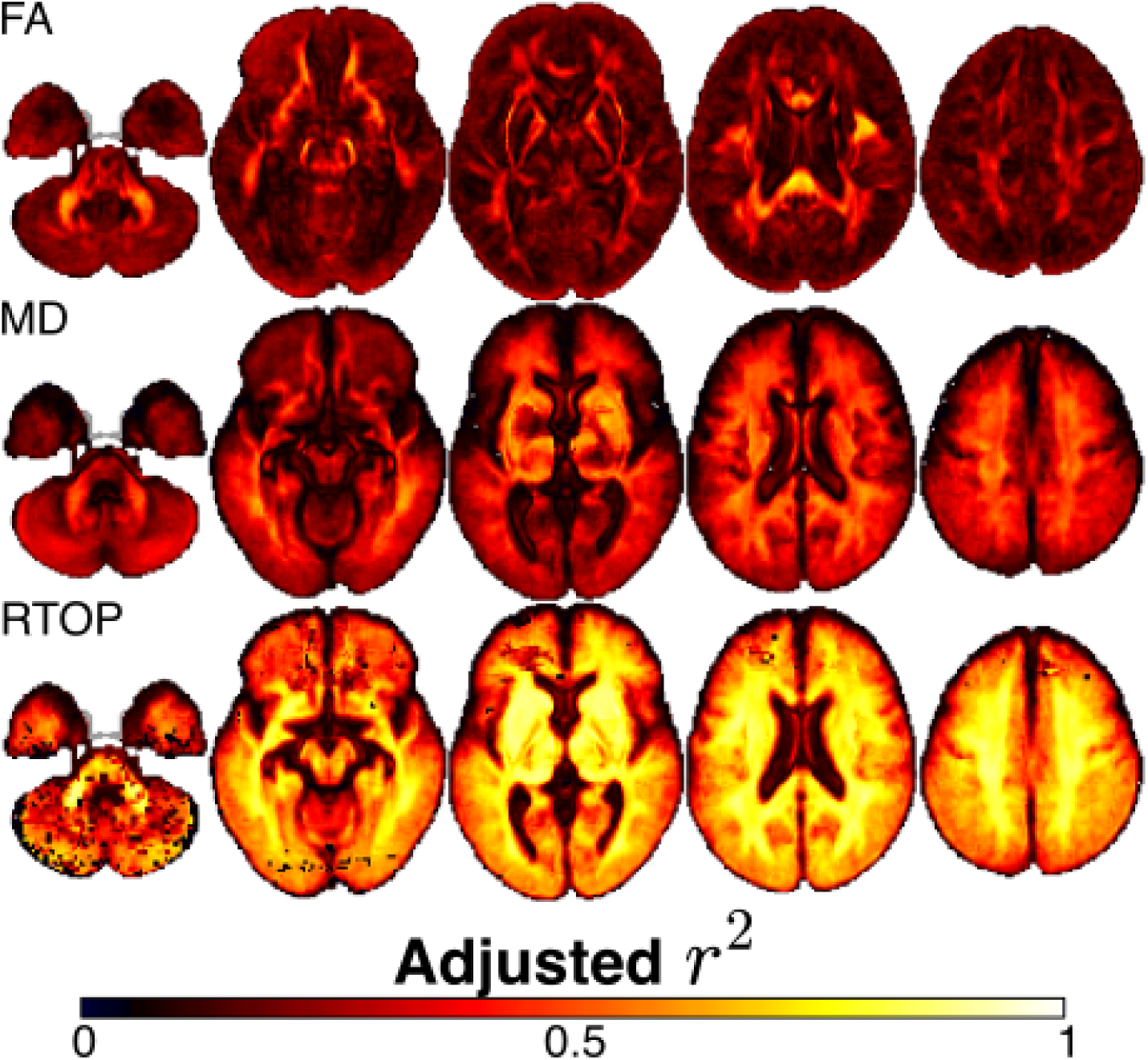
The adjusted for the model fits displayed in Figure 7 are shown for FA, MD and RTOP. The adjusted r2 is calculated by R’s lm() function, which penalizes the inclusion of model parameters. Note that the design matrix fit in each of these models is identical, including NDC, scanning site, vendor and gestational age to predict FA, MD or RTOP.

**Supplementary Figure 5.**
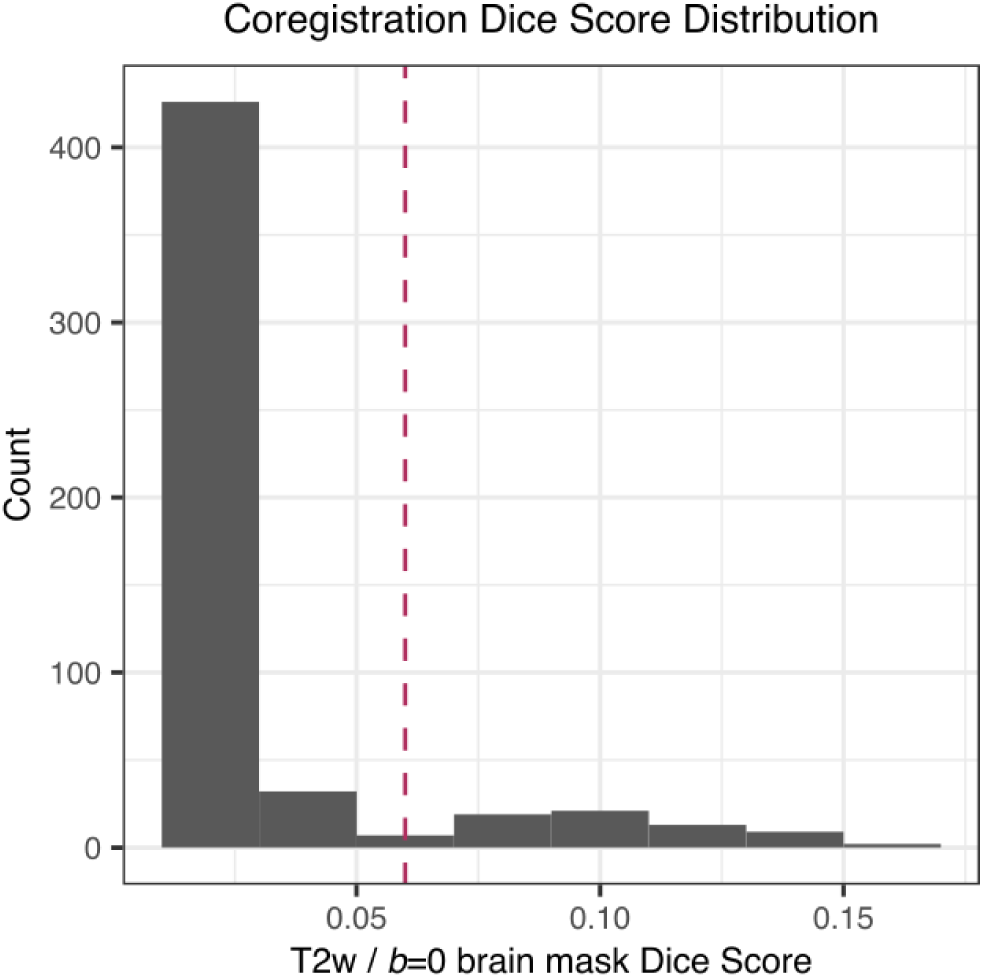
Following coregistration and resampling, a Dice distance was calculated based on the SynthSeg brain masks based on the T2w and mean *b*=0 images from each session. Higher Dice distances indicate lower overlap between the masks and poor coregistration. A vertical dashed line at reflects the cutoff selected for inclusion in the mass univariate analysis in Figure 7.

**Supplementary Figure 6.**
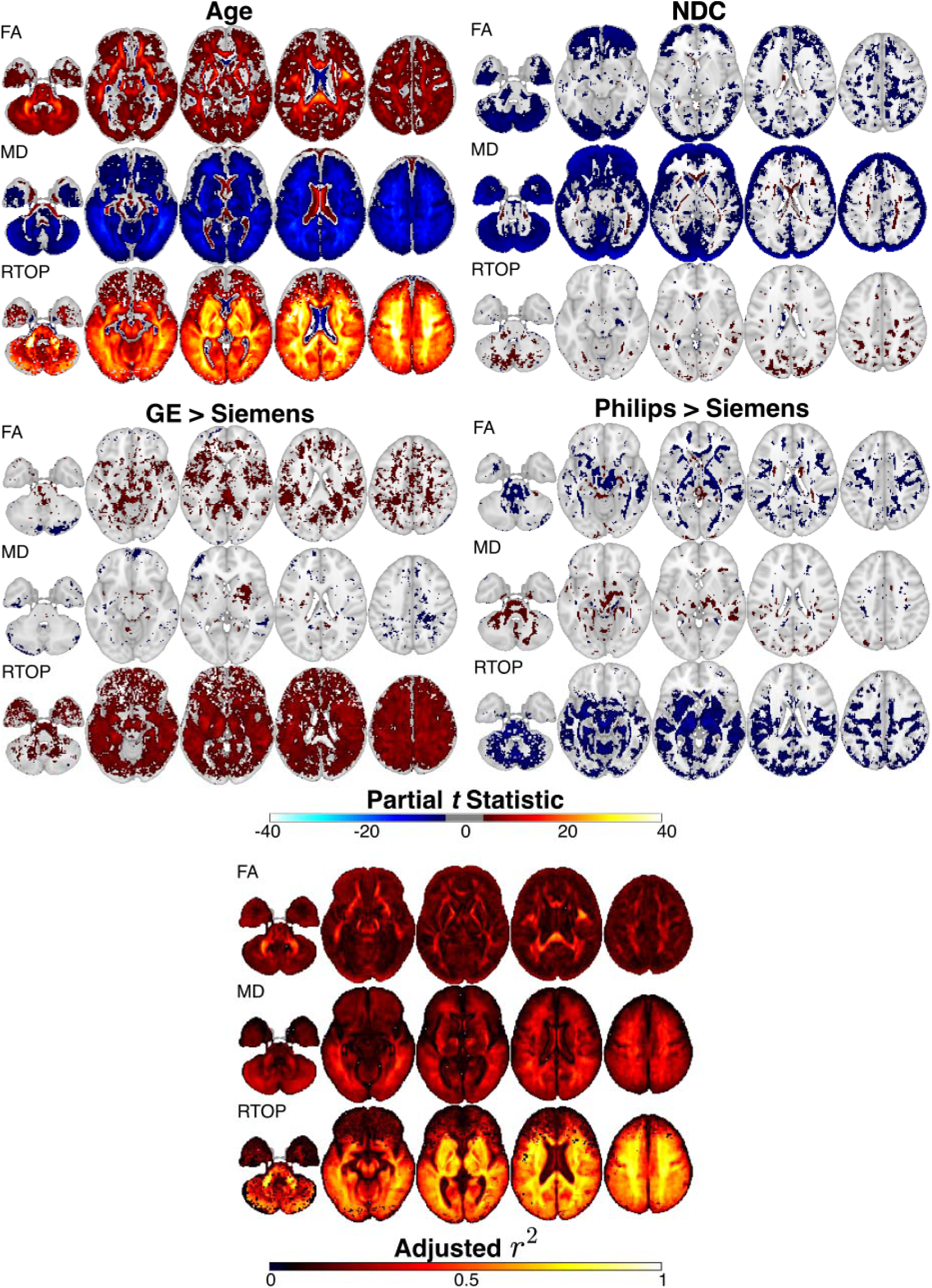
Statistical tests including all data (*n*=529). Including sessions with high coregistration Dice distance reduces overall and specifically impacts the RTOP model fits in the front of the brain.

**Supplementary Figure 7.**
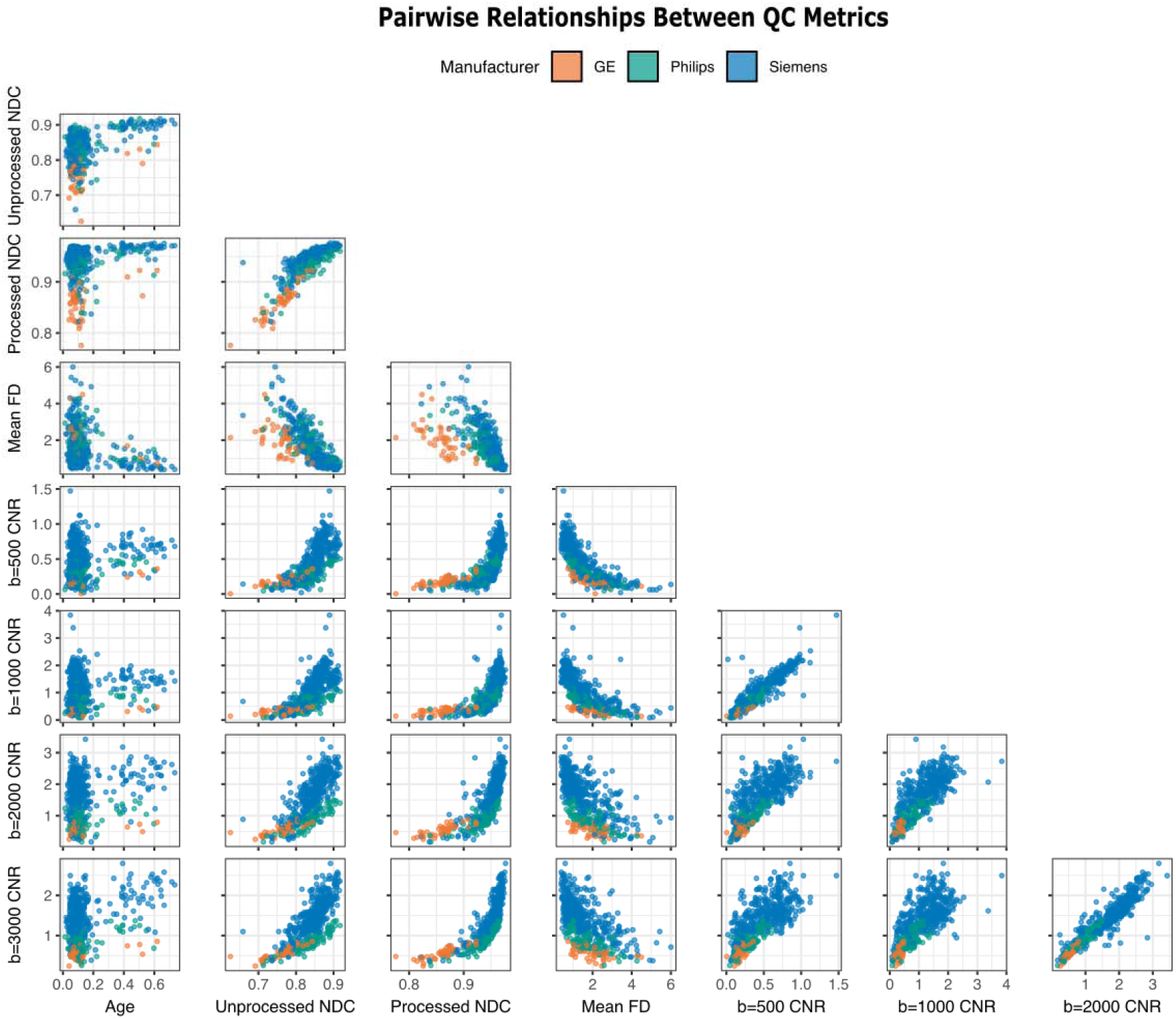
The relationship between age, head motion and automated QC metrics from QSIPrep shows that mean framewise displacement (FD) is highly correlated with both unprocessed and processed NDC. The NDC measures are also highly correlated with CNR, which also correlates with itself at each of the different shells.

**Supplementary Figure 8.**
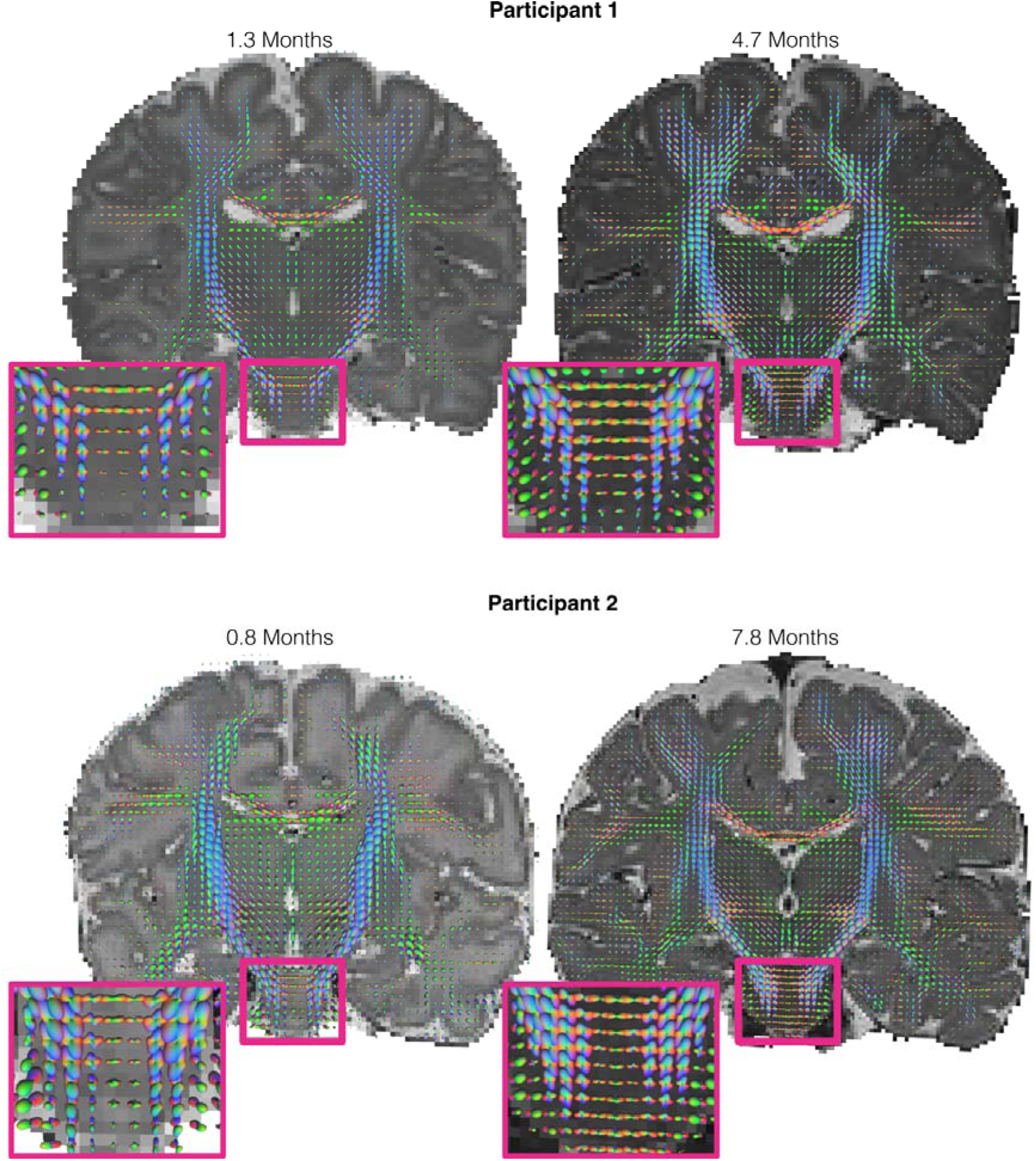
Diffusion ODFs and major white matter bundles plotted for two longitudinal participants. Each row shows an expanded view of the pons (inset) and a coronal slice of the T2w image with dODFs plotted as scaled 3D glyphs. Size and brightness of the glyphs are scaled by anisotropy. The ses-V02 dODFs are less anisotropic as expected, with ses-V03 dODF glyphs both larger and brighter. Notably, crossings are visible in the dODFs in the pons in both sessions for both participants in the inset.

## Supplementary Methods (Automated Boilerplate)

Preprocessing was performed using *QSIPrep* 1.0.1 (Cieslak et al., 2021), which is based on *Nipype* 1.9.1 [(Gorgolewski et al., 2011); RRID:SCR_002502]. Many internal operations of *QSIPrep* use *Nilearn* 0.10.1 (Abraham et al., 2014) and *Dipy* (Garyfallidis et al., 2014).

The T2-weighted anatomical reference image was reoriented into AC-PC alignment via a 6-DOF transform extracted from a full affine registration to the MNIInfant template appropriate for the participant’s age. A full nonlinear registration to the template from AC-PC space was estimated via symmetric nonlinear registration (SyN) using antsRegistration. Brain extraction was performed on the T2w image using *SynthStrip* (Hoopes et al., 2022) and automated segmentation was performed using *SynthSeg* (Billot et al., 2023) from *FreeSurfer* version 7.3.1.

Initial denoising steps were applied to the AP and PA acquisitions *separately.* DWI data were denoised using the Marchenko-Pastur PCA method implemented in *dwidenoise* (Tournier et al. 2019; Veraart et al. 2016; Veraart, Fieremans, and Novikov 2016; Cordero-Grande et al. 2019) with a window size of 3 voxels. After MP-PCA, Gibbs ringing was removed using *TORTOISE* (H.-H. Lee et al., 2021). Any images with a b-value less than 100 s/mm^2^ were treated as a *b*=0 image. The mean intensity of the DWI series was adjusted so all the mean intensity of the *b*=0 images matched across the AP and PA scans. B1-(receive-field) inhomogeneity correction was not included during preprocessing because this artifact was corrected for on the scanner (i.e., Prescan Normalize, PURE, and CLEAR for Siemens, GE, and Philips, respectively). The AP and PA scans were then concatenated for processing with *eddy*.

*FSL* (version 6.0.7.8)’s *eddy* was used for head motion correction and *eddy* current correction (Andersson et al., 2016). *eddy* was configured with a *q*-space smoothing factor (a.k.a the “fudge factor”) of 10, a total of 5 iterations, and 1000 voxels used to estimate hyperparameters. The FWHM of the first iteration in *eddy* was increased to allow for larger head motion and potential large differences in head positioning between the AP and PA scans. A quadratic first level model was used to characterize *eddy* current-related spatial distortion. *q*-space coordinates were forcefully assigned to shells. Field offset was attempted to be separated from subject movement. Shells were aligned post-*eddy*. *eddy*’s outlier replacement was run (Andersson et al., 2016). Data were grouped by slice, only including values from slices determined to contain at least 250 intracerebral voxels. Groups deviating by more than 4 standard deviations from the prediction had their data replaced with imputed values.

Data was collected in a pair of scans with reversed phase-encoding directions, resulting in scans with distortions going in opposite directions. Distortion correction was performed in two stages. In the first stage, *FSL*’s *TOPUP* (Andersson et al., 2003) was used to estimate a susceptibility-induced off-resonance field based on *b*=0 images extracted from both DWI series (with reversed phase encoding directions). The *TOPUP*-estimated fieldmap was incorporated into the eddy current and head motion correction interpolation. Dynamic susceptibility distortion correction was applied with 10 iterations, lambda=10.00 and spline knot-spacing of 10.00 mm (Andersson et al., 2018). Interpolation after head motion and initial susceptibility distortion correction was performed using the jac method.

*DRBUDDI* (Irfanoglu et al., 2015), part of the *TORTOISE* (Irfanoglu et al., 2025) software package, was used to perform a second stage of distortion correction. *DRBUDDI* was run on data output by *eddy*: motion-corrected DWI series split by their phase encoding directions. Specifically, tensor-based synthetic *b*=0 and fractional anisotropy images are computed separately for each phase encoding direction. These are then used together with the T2-weighted structural image in a multimodal registration to estimate the residual susceptibility-induced off-resonance field not detected by TOPUP. Signal intensity was adjusted in the final interpolated images using a method similar to *eddy*’s Least Squares Reconstruction (LSR).

Several confounding time-series were calculated based on the preprocessed DWI: framewise displacement (FD) using the implementation in *Nipype* (following the definitions by (Power et al., 2014)) was calculated based on the motion parameters estimated by *eddy*. The head-motion estimates calculated in the correction step were also placed within the corresponding confounds file. Note that estimation of translation in the phase encoding axis has high uncertainty since that particular aspect of head motion is difficult to distinguish from the effect of a constant (mean) eddy-current field. Slicewise cross correlation was also calculated. QC metrics included in Richie-Halford et al. (2022) such as NDC and coherence index, were calculated along with some new additions like CNR0-4 from *eddy* and DWI Contrast from DSI Studio. The DWI time-series were resampled to AC-PC orientation, generating a *preprocessed DWI run in AC-PC space* with 1.7mm isotropic voxels.

## Supplementary Files: TOPUP and eddy settings

~~~
{
"flm": "quadratic",
"slm": "none",
"fep": false,
"interp": "spline",
"nvoxhp": 1000,
"fudge_factor": 10,
"dont_sep_offs_move": false,
"dont_peas": false,
"niter": 5,
"method": "jac",
"repol": true,
"num_threads": 1,
"is_shelled": true,
"use_cuda": false,
"cnr_maps": true,
"residuals": false,
"output_type": "NIFTI_GZ",
"estimate_move_by_susceptibility": true,
"args": ""
}
~~~

## Notes

### Summary of Updates

Conflict of interest section updated; sentence added to clarify data collection

### Funding

- National Institutes of Health: U01DA055371, U24DA055330, U01DA055350, U01DA055365, U01DA055370, U01DA055362

